# YAP promotes cell-autonomous immune responses to tackle intracellular *Staphylococcus aureus in vitro*

**DOI:** 10.1101/2022.05.17.492111

**Authors:** Caire Robin, Audoux Estelle, Thomas Mireille, Dalix Elisa, Peyron Aurélien, Rodriguez Killian, Dickerscheit Yann, Marotte Hubert, Vandenesch François, Laurent Frédéric, Josse Jérôme, Paul. O Verhoeven

**Affiliations:** CIRI, Centre International de Recherche en Infectiologie, GIMAP Team, Univ Lyon, Univ St-Etienne, INSERM U1111, CNRS UMR5308, ENS de Lyon, Université Claude Bernard Lyon 1, St-Etienne, France; SAINBIOSE, U1059-INSERM, Université de Lyon, St-Etienne, France; CIRI, Centre International de Recherche en Infectiologie, Team Staphylococcal Pathogenesis, Univ Lyon, INSERM U1111, CNRS UMR5308, ENS de Lyon, Université Claude Bernard Lyon 1, Lyon, France; Department of Bacteriology, Institute for infectious Agents, Hospices Civiles de Lyon, Lyon, France; Department of Infectious Agents and Hygiene, University Hospital of St-Etienne, St-Etienne, France

**Keywords:** YAP, *Staphylococcus aureus*, autophagy, lysosome, inflammation, C3 exoenzyme, EDIN, cell-autonomous immunity, host response genes

## Abstract

Transcriptional cofactors YAP/TAZ have recently been found to support autophagy and inflammation, which are part of cell autonomous immunity and are critical in antibacterial defense. Here, we studied the role of YAP against *Staphylococcus aureus* using CRISPR/Cas9-mutated HEK293 cells and a primary cell-based organoid model. We found that *S. aureus* infection increases YAP transcriptional activity, which is required to reduce intracellular *S. aureus* replication. A 770-gene targeted transcriptomic analysis revealed that YAP upregulates genes involved in autophagy/lysosome and inflammation pathways in both infected and uninfected conditions. The YAP/TEAD transcriptional activity promotes autophagic flux and lysosomal acidification, which are important for defense against intracellular *S. aureus*. Furthermore, the staphylococcal toxin C3 exoenzyme EDIN-B was found effective in preventing YAP-mediated cell-autonomous immune response. This study provides new insights on the anti-*S. aureus* activity of YAP, which could be conserved for defense against other intracellular bacteria.

**GRAPHICAL ABSTRACT:** 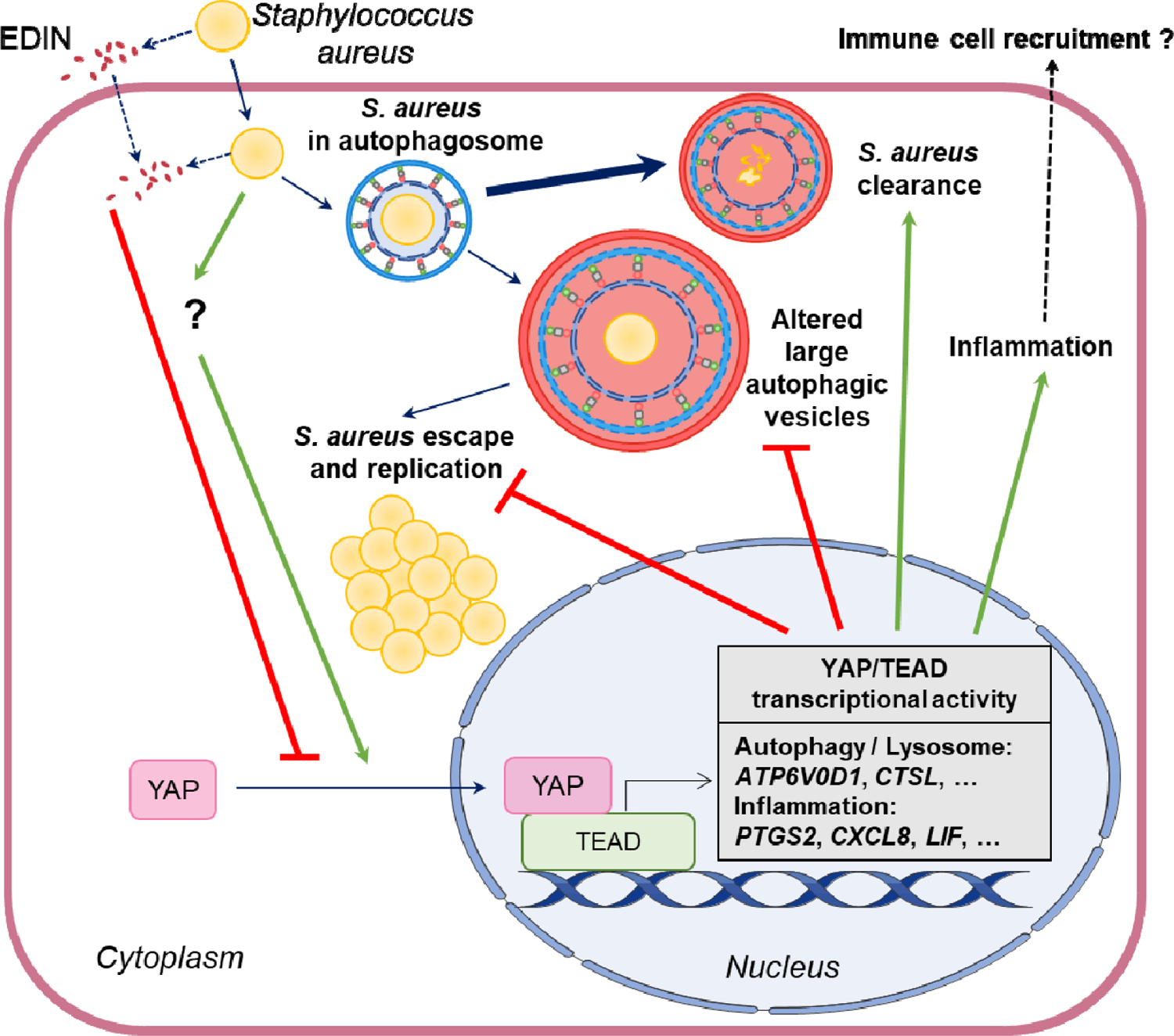

## INTRODUCTION

Yes-associated protein (YAP) and transcriptional co-activator with PDZ-binding motif (TAZ) are transcriptional co-factors involved in many basic cellular functions. YAP and TAZ could interact with TEA domain transcription factor (TEAD), their main transcriptional partner, to elicit target gene expression ^1, 2^. This interaction occurs through the TEAD-binding domain (TBD) of YAP, which is highly conserved throughout evolution ^3, 4^. The Hippo pathway was the first described mechanism for YAP/TAZ phosphorylation that leads to its cytoplasmic retention or proteasomal degradation ^5, 6^. Additionally, YAP/TAZ act as major mechanotransducers that integrate mechanical *stimuli* into transcriptional responses ^7^. The subcellular localization and nuclear translocation of YAP are regulated by the Rho family of GTPases and actin tension^7–9^. At low cell density, YAP exists in the nucleus and is transcriptionally active, whereas at high cell density, it remains in the cytoplasm ^8^. The YAP/TAZ transcriptional program has been extensively studied in cancer research because it promotes cancer cell survival, proliferation, and invasiveness ^10^. Growing evidence suggests that YAP/TAZ are inflammation-responsive and promote inflammation, as well as immune pro-inflammatory cell differentiation ^11–13^. Recent studies have highlighted the role of YAP/TAZ in autophagy through the transcription of genes encoding proteins involved in the formation autophagosomes or their fusion with lysosomes ^14, 15^. Autophagy against intracellular pathogens (formerly called xenophagy) is used by virtually any cell type. Autophagy and inflammation are conserved cell-autonomous responses that restrict infection and increase specialized immune cell recruitment for pathogen clearance ^16, 17^. Despite its involvement in autophagy and inflammation, the modulation and role of YAP during bacterial infections remain poorly investigated, and the findings are somewhat controversial. *Helicobacter pylori* infection in gastric cells (in vitro) leads to YAP transcriptional activation and inflammation (increased IL-1B expression), which, in turn, promotes tumorigenesis ^18^. YAP transcriptional activity in B cells has been found to promote inflammasome activation and likely contribute to defense against *Salmonella* infection in vitro ^19^. In a mouse model of pneumonia due to *Streptococcus pneumoniae*, alveolar cells exhibited increased YAP/TAZ activity, which is important for tissue healing as well as reducing NF-κB activity ^20^. In *C. elegans* and mice, YAP is required to control intestinal infection by *Pseudomonas aeruginosa* and *Salmonella typhimurium* ^21^. In contrast, Yorkie (YAP homolog in *Drosophila melanogaster*) transcriptional activity was found to inhibit the production of anti-microbial peptides by inhibiting NF-κB activity and fostering infection with gram-positive bacteria ^22^. In addition, indirect observations could link YAP and bacterial infections. Indeed, C3 exoenzyme ADP-ribosyltransferase, a bacterial toxin secreted by *Clostridium botulinum*, is known to be a highly specific RhoA inhibitor ^23^. This commercially available toxin is commonly used to inhibit YAP activity *in vitro* ^7^. It is also noteworthy that many intracellular bacterial species can produce C3-like and other toxins that are potent RhoA inhibitors ^24, 25^. For instance, epidermal cell differentiation inhibitors (EDINs) produced by *Staphylococcus aureus* belong to the *Clostridium botulinum* C3 exoenzyme family of bacterial ADP-ribosyltransferases ^26^. The EDIN-B-expressing *S. aureus* clone ST80-MRSA-IV was found to inhibit RhoA activity *in vitro* ^27^. In humans, the prevalence of *edin*-positive *S. aureus* strains is associated with deep-seated infections of soft tissues, suggesting that EDINs increase the virulence of *S. aureus in vivo* ^28^. Despite the strong ability of the C3 exoenzyme to inhibit YAP transcriptional activity, whether the intracellular production of C3 exoenzymes, such as EDINs, could foster *S. aureus* infection through YAP inhibition remains unknown.

*Staphylococcus aureus* is both a commensal and a life-threatening human pathogen responsible for various infections, such as soft skin tissue infections, bacteremia, endocarditis, and osteoarticular infections ^29^. It is widely recognized as a facultative intracellular bacterium capable of triggering its internalization inside non-professional phagocytic cells (NPPCs) by interacting with different host cell receptors ^30^. Inside the host cell, *S. aureus* has been found to be engulfed in autophagosomes by selective autophagy involving cargo receptor proteins, such as sequestosome 1 (SQSTM1/P62), restricting intracellular *S. aureus* ^31^. Autophagy has been shown to be a critical mechanism in defense *against S. aureus* infection in mice and zebrafish ^32, 33^.

In this study, we investigated the potential antibacterial role of the YAP/TEAD transcriptional program using *S. aureus* infection in HEK293 cells and synovial organoid-based models. We showed that YAP/TEAD transcriptional activity is involved in xenophagy because it enhances autophagic flux to promote *S. aureus* clearance. Further, we showed that YAP mediates the expression of host response genes that are known to be important for clearing bacterial infections. In addition, we demonstrated that EDIN-B-producing *S. aureus* prevents YAP/TEAD transcriptional activity and fosters intracellular bacterial replication.

## RESULTS

### *Staphylococcus aureus* infection elicits YAP transcriptional activity prevented by the expression of the *edin*B gene

In this study, we used a lysostaphin (a non-cell permeable bacteriocin active against *S. aureus*) protection assay-based model ^34^ to focus on intracellular bacteria and avoid uncontrolled extracellular bacterial replication.

To investigate YAP signaling in response to *S. aureus* infection, we first used the HG001 *S. aureus* strain (that lacks *edin* genes) in HEK293 cells at different cell densities. At high cell density (HD), YAP was mainly cytoplasmic, as expected (**Figure 1A**). In this *scenario*, *S. aureus* induced an increase in YAP nuclear mean fluorescence intensity (MFI) but not in cytoplasmic MFI, resulting in an increase in the YAP nuclear cytoplasmic (NC) ratio at 7 h post infection (hpi) (**Figure 1A-D**). In contrast, at low cell density (LD) (i.e., when cells are completely isolated from each other, and YAP is exclusively localized in the nucleus) YAP remained localized in the nucleus upon *S. aureus* infection (**S. Figure 1A**). Immunoblotting showed that *S. aureus* did not change YAP and TAZ total protein levels at medium cell density (MD) (i.e., when cells formed few contacts and YAP was mainly localized in the nucleus) (**S. Figure 1B-D**). In MD, neither the activity of TEAD nor the expression of cysteine-rich inducer 61 (*CYR61*), which is a YAP/TAZ/TEAD target gene, was modified upon *S. aureus* infection (**S. Figure 1E-G**). Thus, *S. aureus* HG001 strain infection was found to trigger YAP nuclear translocation but did not enhance YAP signaling when it was already active. We then tested whether the C3 exoenzyme EDIN-B secreted by the *S. aureus* ST80-MRSA-IV strain could prevent YAP activation. Given that the *edin*B-encoded C3 exoenzyme is a membrane non-permeable toxin ^35^, cells were incubated with *S. aureus* culture supernatants for 24 h to allow the toxin to enter cells. We found that the culture supernatant of the ST80 wild-type (WT) strain reduced the nuclear and cytoplasmic localization of YAP, resulting in a decrease in the YAP NC ratio in cells at HD (**S. Figure 2A-D**) and an inhibition of TEAD transcriptional activity at LD (**Figure 1F**). In contrast, the culture supernatant of the *edin*B-deleted ST80-MRSA-IV strain (ST80 *edin*B) had no effect on YAP localization and TEAD activity (**Figure 1F and S. Figure 2A-D**). Together, these results demonstrate that *S. aureus* EDIN-B toxin is highly effective in inhibiting YAP/TEAD activity. *Staphylococcus aureus* has been shown to be more efficient in delivering EDIN-B directly into the host cell after internalization ^35^. Consequently, we tested whether infection with the ST80 WT and ST80 Δ*edin*B strains modulates YAP subcellular localization and transcriptional activity. As expected, the ST80 *edin*B strain was found to enhance YAP nuclear intensity and decrease YAP cytoplasmic intensity, resulting in a strong increase in the YAP NC ratio (**Figure 1A-D**) at 7 hpi in HD cells. In contrast, the EDIN-B-expressing ST80 WT strain was found to reduce YAP nuclear MFI and NC ratio compared to ST80 Δ*edin*B at 7 hpi in HD cells as well as YAP cytoplasmic and nuclear MFI compared to the control cells (**Figure 1A-D**). In addition, ST80 Δ*edin*B was found to increase TEAD transcriptional activity as soon as 3 hpi in HD cells, whereas it was not the case with the ST80 WT strain (**Figure 1E**). These results demonstrated that the *S. aureus* infection (but not the *S. aureus* supernatant) caused an increase in YAP nuclear localization and YAP/TEAD transcriptional activity *in vitro*. Interestingly, the EDIN-B-expressing ST80 WT strain as well as the EDIN-B toxin alone were found to be effective in preventing or decreasing YAP activity.

**Figure 1.**
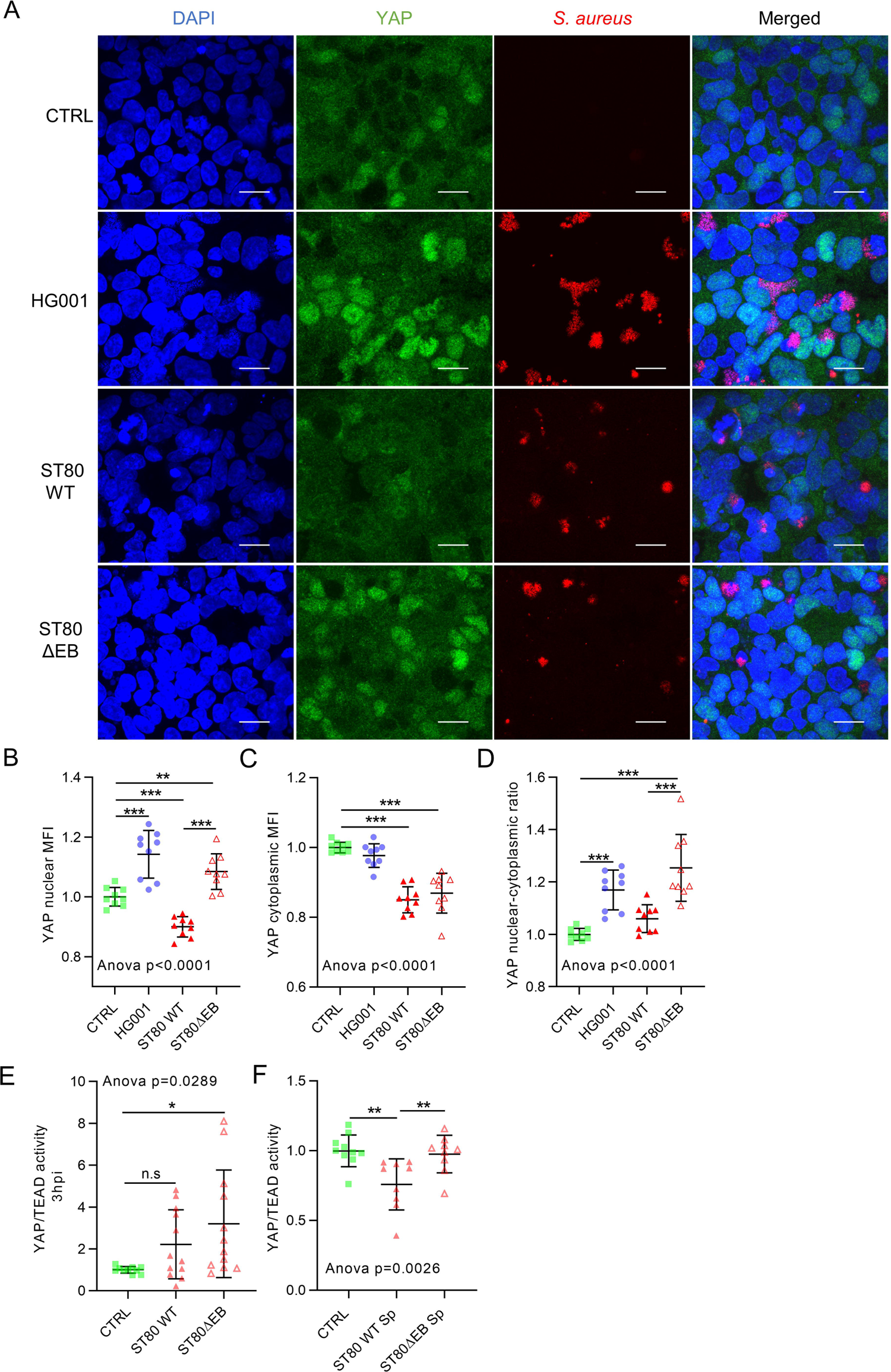
*Staphylococcus aureus* toxin EDIN-B prevented YAP activation in HEK293 cells HEK293 cells were cultured at high. (A-E) or low density (F). HG001 or ST80 *S. aureus* infection was at a multiplicity of infection of 10 for 7 h (A-D) or 3 h (E). *S. aureus* were allowed to contact for 2 h with the cells, and lysostaphin was added at 10 µg/mL for the rest of the experiments to avoid extracellular *S. aureus* multiplication. A: Confocal representative z-stack max intensity projection images of YAP (immunolabeling, green), DAPI (nucleus, blue), *S. aureus* (DsRed, red), and merged. Scale bar: 20 µm. B-D: Quantification of YAP nuclear mean fluorescence intensity (MFI) (B), YAP cytoplasmic MFI (C), and YAP nuclear cytoplasmic ratio (D) of HG001-infected cells. E-F: luciferase reporter assay of TEAD transcription factor activity (8xGTIIC) for ST80 *S. aureus* infection (E) or ST80 strain supernatant treatment for 24 h (F). Results were expressed as fold change vs. control group and presented as individual values with mean ± SD, representing three independent experiments. CTRL: control; WT: wild-type; ST80ΔEB: ST80 EDIN-B-deleted strain; Sp: supernatant. ANOVA test with false discovery rate (FDR) correction for multiple comparisons post hoc tests: * p<0.05, ** p<0.01, *** p<0.001.

### YAP transcriptional activity is required to limit the intracellular replication of *S. aureus*

As YAP is activated during infection, we investigated whether YAP/TEAD transcriptional activity was needed to fight *S. aureus in vitro*. We used WT and YAP-deleted (YAP^-/-^) HEK293 cells generated using the CRISPR-Cas9 technique ^36^. YAP knockout was confirmed by immunoblotting, and the absence of YAP transcriptional activity was confirmed by a decrease in *CYR61* expression by RT-qPCR (**S. Figure 1B, C, and G**). Interestingly, YAP knockout decreased TAZ total protein levels (**S. Figure 1B and D**), which can contribute to decreased TEAD transcriptional activity. To specifically investigate the role of YAP/TEAD activity, we engineered HEK293 cells with a heterozygote mutation of YAP within its TEAD-binding domain (YAPΔTEAD^-/+^) that resulted in the substitution of four amino acids (**Figure 2A**) critical for binding to TEAD ^4^. In LD cells, YAP/TEAD activity was strongly decreased in YAP TEAD^-/+^ cells compared to WT cells (**Figure 2B**). Subsequent experiments were performed at MD to have a robust basal activity of YAP in WT cells, compared to YAP-mutated cells (i.e., YAP^-/-^ and YAPΔTEAD^-/+^ cells). Using the DsRed-expressing *S. aureus* HG001 strain, we first observed by confocal microscopy that *S. aureus* was able to replicate in WT cells between 3 and 7 hpi. Strikingly, *S. aureus* intracellular replication was more pronounced in both YAP^-/-^ and YAP TEAD^-/+^ cells, with the presence of heavily infected cells (**Figure 2C and D**). These results were confirmed by quantifying intracellular *S. aureus* loads on agar plates (**Figure 2E**). Interestingly, in WT cells, the intracellular replication of the ST80 WT strain was more pronounced than that of the ST80 ΔEB strain, showing that EDIN-B expression was an advantage for *S. aureus* intracellular replication *in vitro* (**Figure 2C, F**). Thus, YAP/TEAD activity and EDIN-B expression exerted opposite effects by restricting and enhancing the intracellular replication of *S. aureus*, respectively.

**Figure 2.**
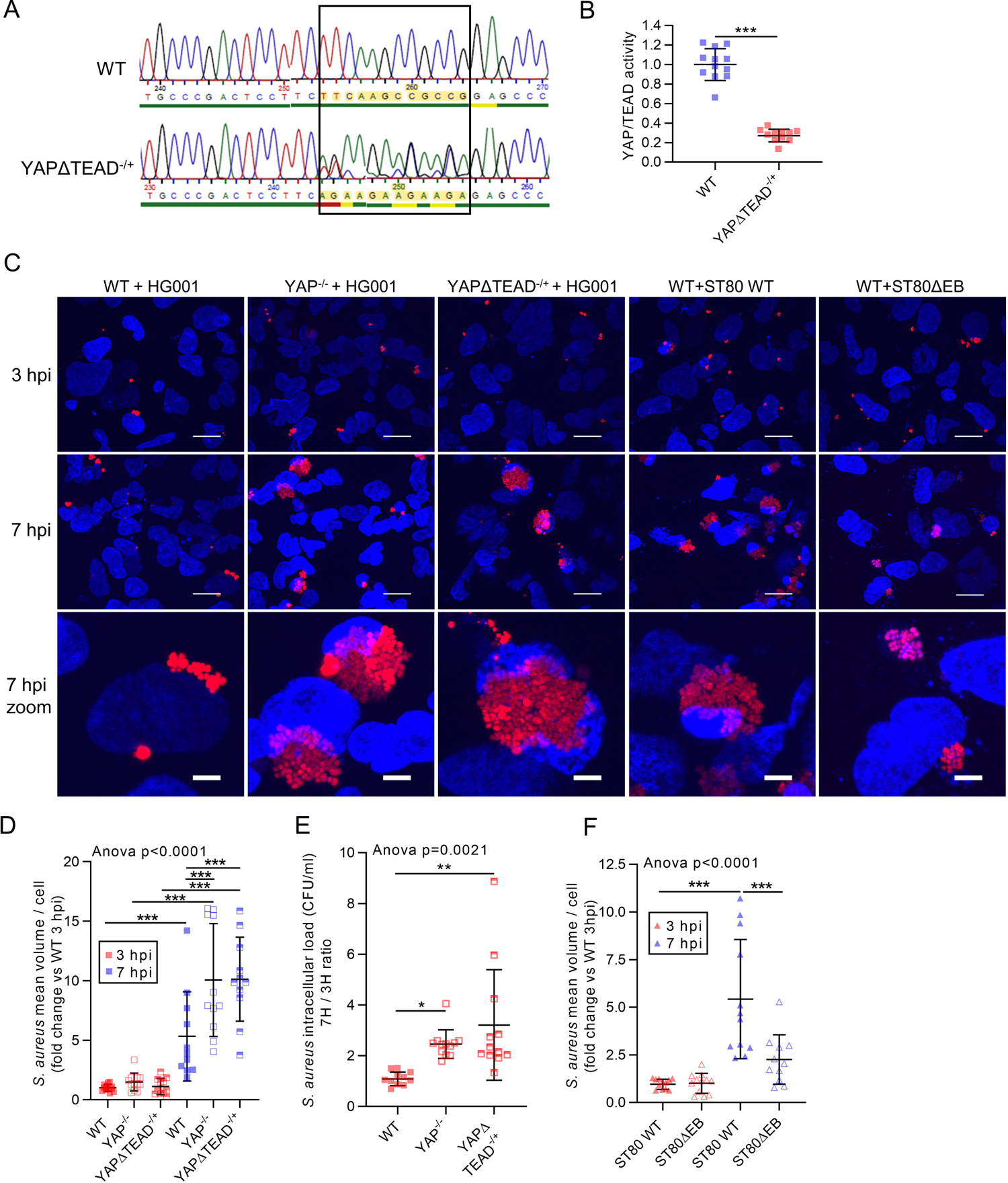
YAP transcriptional activity inhibits intracellular *Staphylococcus aureus* replication. HEK293 cells were cultured at low (B) or medium density (C-F). HG001 or ST80 *S. aureus* infection was at a multiplicity of infection of 1 for 3 or 7 h, as indicated. *Staphylococcus aureus* were allowed to contact cells for 2 h, and lysostaphin was added at 10 µg/mL for the remaining experiments to avoid extracellular *S. aureus* multiplication. A: Electrophorogram of WT and YAPΔTEAD^-/+^ cells showing TTCAAGCCGCCG replacement by AGAAGAAGAAGA. B: Luciferase reporter assay of TEAD transcription factor activity (8xGTIIC) for WT and YAPΔTEAD^-/+^ cells; C: Representative confocal z-stack max intensity projection images of live cells labeled with DAPI (nucleus, blue) and infected with DsRed *S. aureus* HG001 or ST80 strains as indicated (red); scale bar 20 µm and 5 µm for zoomed image D: Microscopy quantification of intracellular HG001 mean volume per cell. E: Quantification of HG001 colony forming unit (CFU) per mL on an agar plate at a ratio of 3 hpi/7 hpi. F: Microscopy quantification of the intracellular ST80 strain mean volume per cell. For microscopy quantification (D and F), the total *S. aureus* volume measured in the field was divided by the number of nuclei in the same field. Results were expressed as fold change vs. control group and presented as individual values with mean ± SD, representing three independent experiments. WT: wild type; ST80ΔEB: ST80 EDIN-B-deleted ST80 strain; CFU: colony-forming unit. Unpaired t-test (B) or Analysis of variance (ANOVA) test with false discover rate (FDR) correction for multiple comparison post hoc tests: * p<0.05, ** p<0.01, *** p<0.001.

### YAP is critical to promote the expression of host response genes usually induced by *S. aureus* infection

To understand why YAP transcriptional activity was important in inhibiting *S. aureus* intracellular replication, we analyzed the expression of 770 genes involved in host response in control or HG001-infected WT or YAP^-/-^ cells at 7 hpi and at MD using the nCounter host response panel. Since we showed that *S. aureus* did not increase YAP activity at this cell density, we focused more on the differences between YAP^-/-^ and WT cells in both uninfected and infected conditions.

Striking differences were observed between WT and YAP^-/-^ cells under both uninfected and infected conditions. For instance, 240 genes were downregulated, whereas only 52 were upregulated in YAP^-/-^ infected cells compared to WT infected cells. Most of the downregulated signaling pathways in YAP^-/-^ cells were inflammation-related signaling pathways (e.g., chemokine, interleukin, inflammasome, and prostaglandin signaling pathways) (**Figure 3A**). Upon *S. aureus* infection in WT cells, a pro-inflammatory response profile was induced, whereas in YAP^-/-^ cells, this response was induced but remained at lower levels than that in WT control or infected cells (**Figure 3A**). At the level of individual genes, those encoding pro-inflammatory cytokines and chemokines, such as IL-11, CXCL8, and LIF, were among the most downregulated genes in YAP^-/-^ infected cells compared to WT infected cells (**Figure 3B**). In a few upregulated genes in YAP^-/-^ infected cells compared to WT infected cells, we detected lysosomal genes such as *LAMP1*, *NPC2*, and *GBA* (**Figure 3B**). In YAP^-/-^ cells, we found an upregulation of the lysosome pathway and a downregulation of the autophagic pathway (**Figure 3A**), which are known to reduce the intracellular replication of *S. aureus.* Altogether, these results indicate the involvement of YAP in host response gene expression and its contribution to transcriptional immune response in HEK293 cells, consistent with the gene expression profile induced by *S. aureus* infection.

**Figure 3.**
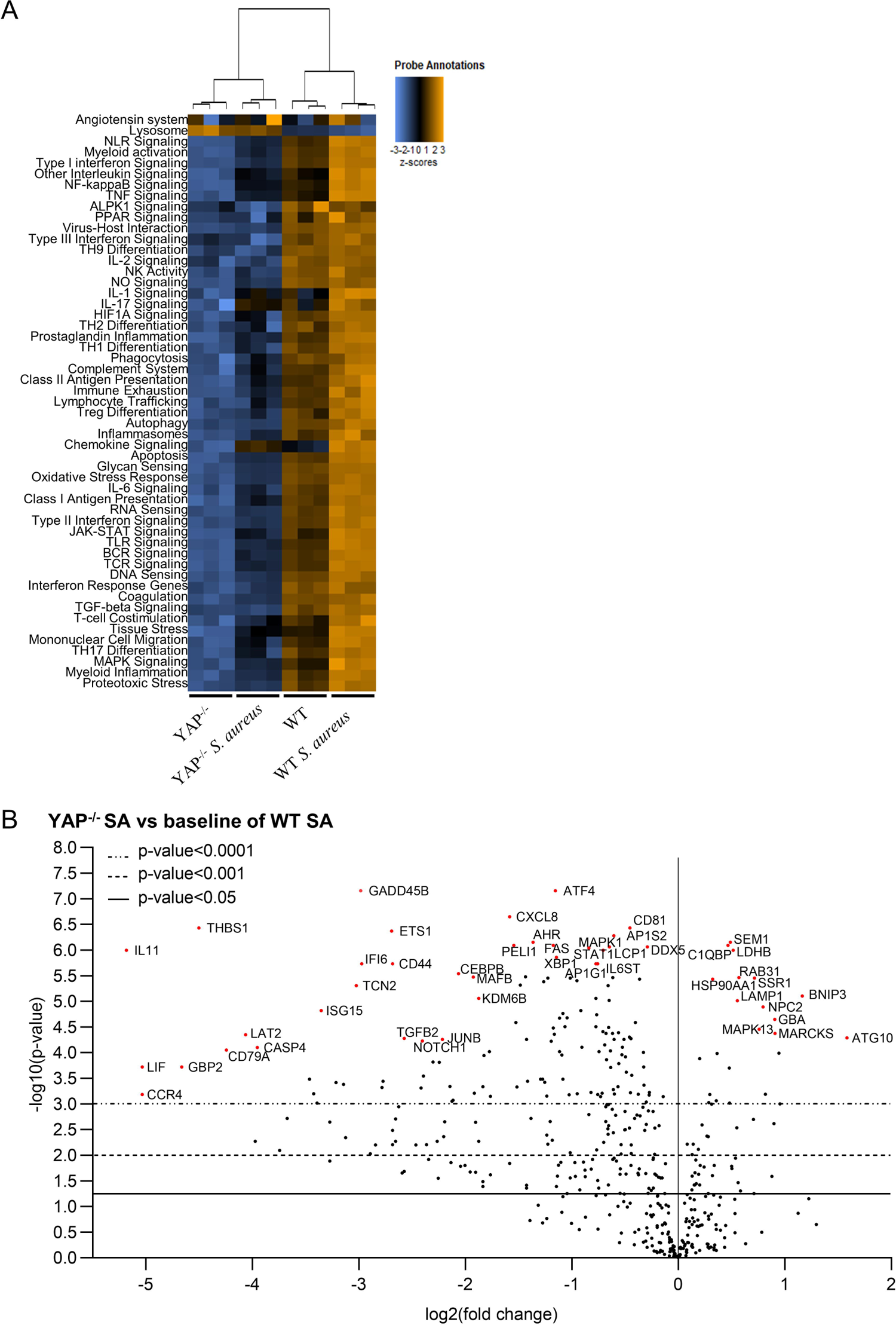
YAP promotes host response gene expression important during *Staphylococcus aureus* infection. HEK293 cells were cultured at medium density and infected with HG001 *S. aureus* strain at a multiplicity of infection of 10 for 7 h. *Staphylococcus aureus* were allowed to contact cells for 2 h with the cells. Subsequently, lysostaphin was added at 10 µg/mL for the remaining experiments to avoid extracellular *S. aureus* multiplication. A: Heat map of nCounter NanoString host response pathways; pathways are listed to the left, the most upregulated pathways are depicted in orange, and the most downregulated pathways are shown in blue; each column corresponds to one sample (n=3 / group). C: Volcano plot representation of differential gene expression in YAP^-/-^ infected group versus the baseline of WT infected group; depicted genes were the most differentially expressed with the combination of a low p-value and a high fold change; p-value was calculated with the NanoString software based on t-test corrected with false discovery rate. WT: Wild type; SA: *S. aureus*.

### YAP/TEAD transcriptional activity regulate autophagic flux and lysosomal acidification

We then decided to focus on the modulation of autophagy and lysosome signaling pathways by YAP activity in non-infected conditions since these processes are critical for defense against intracellular bacteria. In our model, the overall increase in the lysosome signaling pathway in YAP^-/-^ cells was mainly due to an increased expression of genes encoding lysosomal membrane proteins, which could be used as lysosome markers (e.g., *LAMP1*, *NPC2*, and *GBA*) (**Figure 4A**). In contrast, we found decreased expression of several genes related to lysosomal functions. Indeed, we found a decrease in the expression of the *AP1S2* and *AP1G1* genes encoding adaptins that are involved in lysosomal enzyme transport from the trans-Golgi network to lysosomes ^37^. Furthermore, we observed a downregulation in the expression of cathepsin L (*CTSL*) and W and an upregulation in the expression of cathepsin A and Z. A previous study has shown that *CTSL* inhibition leads to LC3-II accumulation and lysosomal enlargement in macrophages ^38^. In addition, we found that *ATP6V0D1*, a gene encoding a subunit of the V-ATPase lysosomal pump critical for lysosomal acidification and autophagy ^39^, was downregulated in YAP^-/-^ cells (**Figure 4A and S**. **Figure 4A**). Interestingly, a chromatin immunoprecipitation assay using next-generation sequencing (ChIP-seq) data from previous reports ^40^ revealed YAP/TAZ/TEAD peaks at active enhancer sites of the *ATP6V0D1*, *ATP6V0A1*, *ATP6V1C1*, and *ATP6V0B* genes ^40^. Thus, this transcriptional profile indicates potential lysosome defects in YAP^-/-^ cells that are modulated by YAP/TEAD transcriptional activity. In addition, several autophagy-related genes, including *MAP1LC3A* (encoding microtubule-associated protein 1 light chain 3 alpha (LC3A) protein), *ATG12* (involved in autophagosome elongation through the LC3-I to LC3-II lipidation), and *ATG13* (involved in autophagosome formation), were downregulated. In addition, we observed an upregulation in *ATG10* (involved in the formation of the ATG5-ATG12-ATG16L elongation complex) that probably compensates for *ATG12* downregulation (**Figure 4A and S**. **Figure 4A**). This profile argued for default autophagosome formation and elongation in YAP^-/-^ cells. Given that YAP/TAZ control the expression of actin-related tension proteins MLC2 and DIAPH1, which are important for autophagosome formation ^14^, we assessed the expression of these two genes; however, MLC2 expression was not detected in HEK293 cells and DIAPH1 expression was similar in HEK293 WT and YAP^-/-^ cells (data not shown).

**Figure 4.**
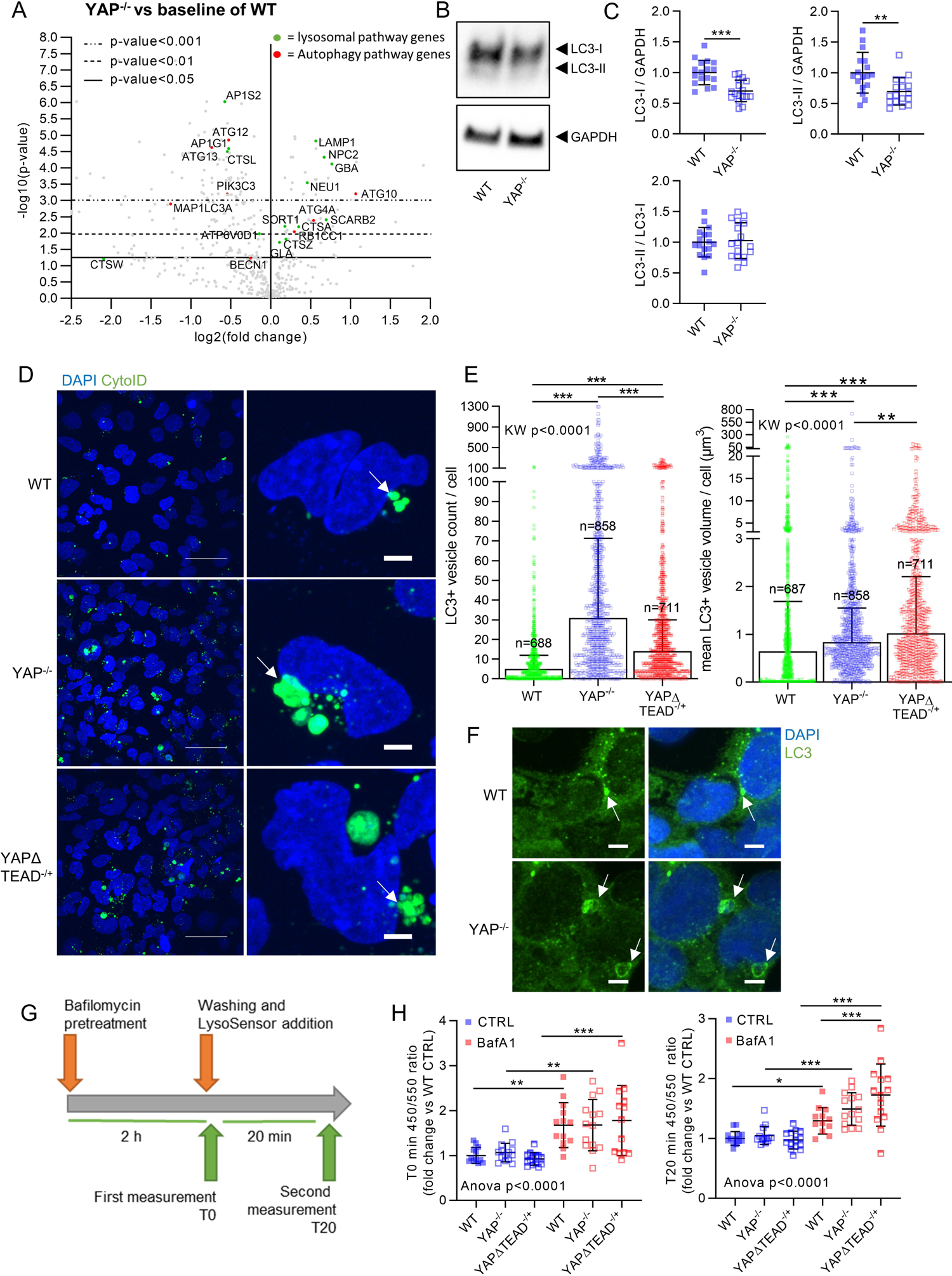
YAP/TEAD activity is involved in autophagic flux regulation *via* lysosomal acidification. HEK293 cells were cultured at medium density and remained uninfected. A: Volcano plot representation of differential gene expression (nCounter NanoString) in YAP^-/-^ control group versus the baseline of WT control group; depicted genes are autophagic (red) or lysosomal (green) pathway genes differentially expressed. B-C: Representative western blot results of LC3A/B -I and -II, and GAPDH (B), with their quantification normalized by GAPDH expression (C). D: Representative confocal z-stack max intensity projection images of live cells labeled with DAPI (nucleus, blue) and CytoID (LC3-II vesicles, green). Scale bar: 20 µm (left) and 5 µm (right). E: corresponding quantification of the LC3-II positive vesicle count or mean vesicle volume per cell, as indicated. Each point represents one cell, the number of analyzed cells per group is indicated; F: Confocal representative z-stack max intensity projection images of LC3 (immunolabeling, green) and DAPI (nucleus, blue); scale bar: 5 µm G: timeline of LysoSensor experiment; bafilomycin pre-treatment was performed for 2 h, cells were washed, and the relative 450/550 nm ratio was assessed immediately or 20 min after LysoSensor addition. H: corresponding results indicating the relative acidity of lysosomes in control or BafA1-treated cells after 0 or 20 min as indicated. Notably, an increase of 450/550 nm ratio is representative of lysosome alkalinization. Results are expressed as fold change vs. control group (only in C and H), and presented as individual values with mean ± SD (C and H) or median with interquartile range (E), representing three independent experiments. WT: wild type; BafA1: Bafilomycin A1. Unpaired t-test (C), Analysis of variance (ANOVA) or Kruskal-Wallis (KW) test with false discovery rate (FDR) correction for multiple comparisons post hoc tests: * p<0.05, ** p<0.01, *** p<0.001.

To confirm the findings of the transcriptional analysis, we monitored autophagic flux in WT, YAP^-/-^, and YAPΔTEAD^-/+^ cells. The immunoblotting assay showed a decrease in the LC3-I total protein level in YAP^-/-^ cells, corroborating the transcriptomic results (**Figure 4BC**). Moreover, the decrease in LC3-I was associated with a decrease in the LC3-II total level, without a change in the LC3-II/LC3-I ratio, suggesting that LC3 lipidation was retained (**Figure 4BC**). To further study autophagy regulation in both models, we used live-cell confocal microscopy and a CytoID probe to label autophagic vesicles in living cells. Autophagic vesicles were more abundant and especially much larger in YAP^-/-^ and YAPΔTEAD^-/+^ cells than in WT cells (**Figure 4DE**). In YAP-mutated cells, these larger autophagic vesicles also appeared misshapen, in contrast to the spherical vesicles observed in WT cells (**Figure 4D**). Similar results were obtained from cells immunolabeled with anti-LC3 antibody (**Figure 4F**), ruling out an artifact due to CytoID. Together, these results reflect autophagic flux reduction resulting in the accumulation of large autophagic vesicles, which suggests an anomaly in the degradative activity of autophagolysosomes. For instance, lysosomal alkalinization is known to induce the accumulation of autophagic vesicles and larger autophagolysosomes ^39^. Since our transcriptomic results indicate possible defects in lysosomal acidification, we tested lysosomal acidity in YAP-mutated cells. No difference was detected in basal conditions between WT, YAP^-/-^, and YAPΔTEAD^-/+^ cells. Bafilomycin A1, an inhibitor of the V-ATPase pump, was effective in inducing lysosomal alkalinization in both cell lines (**Figure 4GH**). However, 20 min after bafilomycin A1 removal, lysosomes reacidification was more efficient in WT cells than in YAP^-/-^ and YAPΔTEAD^-/+^ cells, indicating lysosomal dysfunction in these cells (**Figure 4GH**). These results showed that YAP/TEAD activity promotes the expression of autophagic and lysosomal genes that are important for normal autophagic flux regulation and lysosomal acidification.

### Loss of YAP/TEAD transcriptional activity worsens blockage of autophagic flux induced by *S. aureus* and fosters its escape from autophagic vesicles

Internalized *S. aureus* is known to elicit a strong autophagic response in NPPCs, which is required to clear intracellular *S. aureus* by addressing *S. aureus-*containing autophagosomes to lysosomes. Therefore, we decided to investigate how the alteration of autophagy and lysosome signaling pathways observed in YAP-mutated cells could explain the strong replication of intracellular *S. aureus* in these cells.

Transcriptomic analysis of WT and YAP^-/-^ cells infected with *S. aureus* showed that the expression of specific genes involved in autophagy and lysosome signaling pathways were altered in YAP^-/-^ cells (**S. Figure 3AB**). For instance, *CTSL* expression was still lower in YAP^-/-^ infected cells compare to WT infected cells (**S. Figure 3A-C**).

Live-cell confocal microscopy was used to quantify the number and volume of autophagic vesicles. In WT cells, we observed a strong increase in autophagic vesicle count and volume at 3 hpi, whereas at 7 hpi, the volume of autophagic vesicles continued to rise with no further increase in the vesicle count, which is in favor of the blockage of autophagic flux (**Figure 5A-C**). Interestingly, in WT cells, the colocalization of *S. aureus* with autophagic vesicles was found to decrease between 3 and 7 hpi (**S. Figure 4A and Figure 5D**), which reflects the ability of *S. aureus* to escape from autophagic vesicles (*e.g.*, autophagosomes or autolysosomes). In some of the WT cells, we also observed disrupted *S. aureus* and diffused red fluorescence within autophagic vesicles, indicating that the degradative function of autophagolysosomes can limit the intracellular replication of *S. aureus* in WT cells (**Figure 5A**). In contrast, such a pattern of degradation was not observed in YAP^-/-^ and YAPΔTEAD^-/+^ cells, suggesting that lysosomal degradative function is altered in these cells, which is in accordance with our data of transcriptomic and LysoSensor analyses.

**Figure 5.**
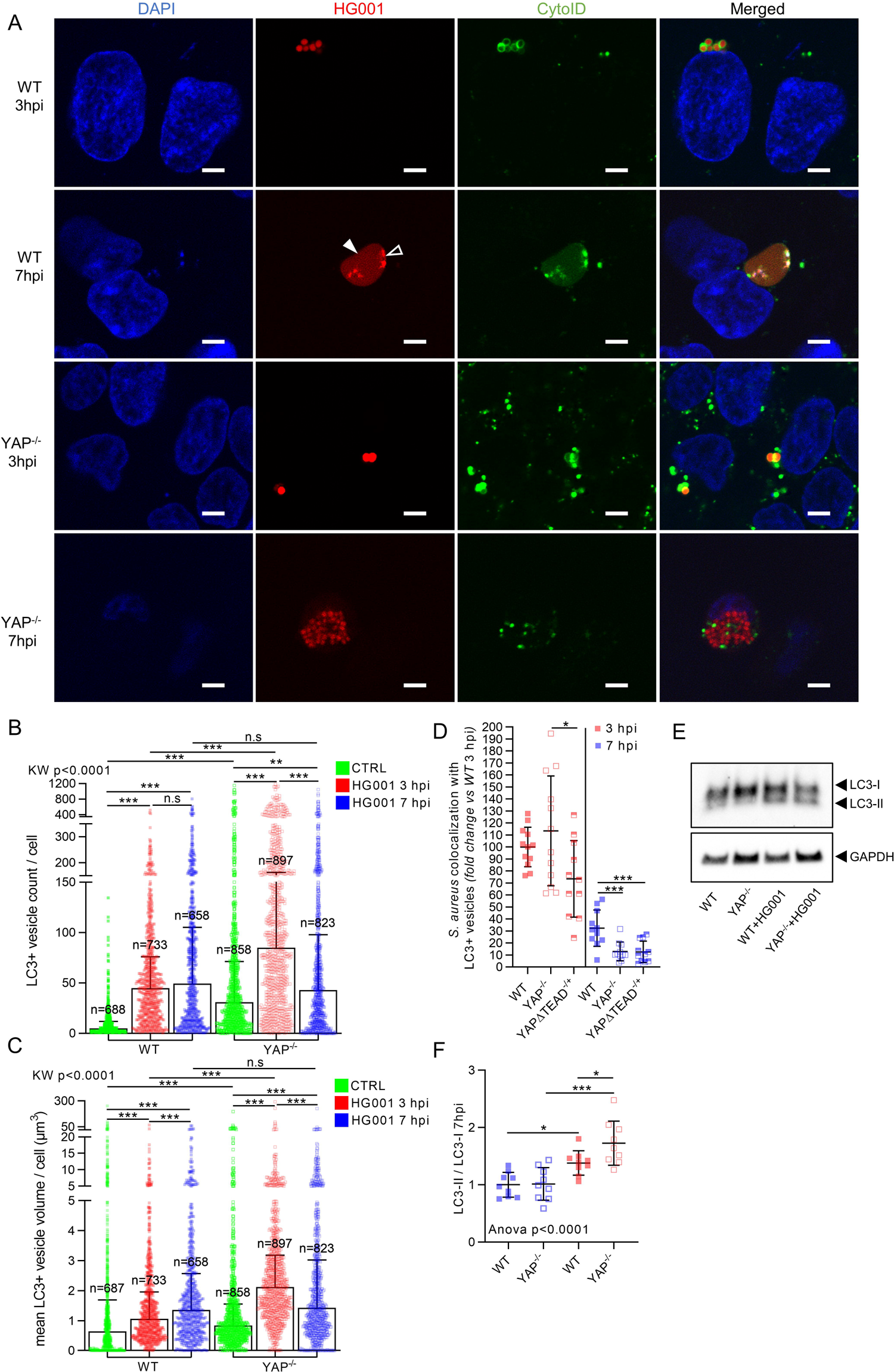
YAP/TEAD transcriptional activity is required to reduce *Staphylococcus aureus* induced autophagic flux blockage. HEK293 cells were cultured at medium density. HG001 *S. aureus* infection was at a multiplicity of infection of 1 for 3 or 7 h, as indicated. *Staphylococcus aureus* were allowed to contact cells for 2 h, and lysostaphin was added at 10 µg/mL for the remaining experiments to avoid extracellular *S. aureus* multiplication. A: Representative confocal (0.5 µm thick z-stack) images of live cells labeled with DAPI (nucleus, blue), CytoID (LC3-II vesicles, green), HG001 (DsRed, red), and merged; white arrowhead: diffused red fluorescence within autophagic vesicles. White empty arrowhead: disrupted *S. aureus.* scale bar: 5 µm. B-C: corresponding quantification of the LC3-II-positive vesicle count (B) or mean volume (C) per cell as indicated. Each point represents one cell. The number of analyzed cells per group is shown. D: Quantification of the relative percentage of colocalization between *S. aureus* and CytoID labelling (LC3-II vesicles). The results are expressed as fold change vs. the WT 3 hpi group set at 100%. E-F: Representative western blot results of LC3-I and II, and GAPDH (E), with their quantification normalized by GAPDH expression (F). Results are expressed as fold change vs. control group (only in D and F) and presented as individual values with mean ± SD (D and F) or median with interquartile range (B and C), representing three independent experiments. WT: Wild type. Analysis of variance (ANOVA) or Kruskal-Wallis (KW) test with false discovery rate (FDR) correction for multiple comparisons post hoc tests: * p<0.05, ** p<0.01, *** p<0.001.

This statement is also supported by the fact that the colocalization of *S. aureus* with autophagic vesicles at 7 hpi in these cells was lower than that observed in WT cells (**Figure 5A, D, and S. Figure 4A**), which reflected *S. aureus* escape from autophagic vesicles. This result did not seem to be due to a defect in autophagy initiation since the autophagic vesicle count increased at 3 hpi compared to non-infected conditions, as it did for WT cells (**Figure 5AB and S. Figure 4AB**). It is also noteworthy that even if the level of colocalization of *S. aureus* with autophagic vesicles was identical to that of WT cells at 3 hpi in YAP-mutated cells, the autophagic vesicles surrounding *S. aureus* were unusually distorted as compared to the spherical vesicles surrounding each individual *S. aureus* bacterium in WT cells (**Figure 5A, D, and S. Figure 4A**). More importantly, the volume of autophagic vesicles strongly increased at 3 hpi in YAP^-/-^ cells than in WT infected cells (**Figure 5A, C**), indicating a further autophagic flux blockage during *S. aureus* infection. In YAPΔTEAD^-/+^ cells, the vesicle volume did not increase further at 3 hpi (**S. Figure 4B, C**), which could be explained by the fact that the vesicle volume in uninfected cells was already higher than that in WT and YAP^-/-^ cells. For YAP-mutated cell lines, vesicle count and volume decreased between 3 and 7 hpi, which seems to be due to the disruption of autophagic vesicles by *S. aureus* that did not colocalize with the spherical vesicles but were surrounded by CytoID-labeled residues (**Figure 5AC and S. Figure 4AC**). Immunoblots showed that the LC3-II/LC3-I ratio was higher during infection in YAP^-/-^ cells compared to WT cells, corroborating that *S. aureus* takes advantage of YAP-deficient cells to further inhibit autophagic flux (**Figure 5EF**).

We then performed similar experiments with WT cells infected with ST80 WT and ST80 Δ*edin*B strains, both of which were found to enhance vesicle count and volume in a very similar manner (**S. Figure 5AC**). Although both ST80 WT and ST80ΔEB strains were found to be highly colocalized in autophagic vesicles at 3 hpi, but the former was able to escape from autophagic vesicles at 7 hpi in contrast to the latter that remained more confined to autophagic vesicles (**S. Figure 5AB**).

Altogether, these results indicate that the alteration of autophagy and lysosome basal functions in YAP-mutated cells enables *S. aureus* to block autophagic flux more efficiently, as soon as 3 hpi, to escape from autophagic vesicles and avoid its clearance into degradative compartments at 7 hpi. These results can explain why *S. aureus* exhibits more pronounced replication in YAP-mutated cells after 3 hpi. In addition, we showed that EDIN-B-expressing *S. aureus* was able to inhibit YAP and was more efficient in escaping autophagy.

### YAP promotes inflammatory response during *S. aureus* infection

Upon bacterial infection, the cell-autonomous immune response of non-specialized immune cells displays antimicrobial mechanisms ^16, 17, 41^. An important part of this response is the activation of molecular signaling pathways that enable the expression of inflammatory mediators to attract specialized immune cells for bacterial clearance.

Our transcriptomic analysis highlighted that most of the differences in gene expression between YAP^-/-^ and WT cells were related to inflammatory signaling pathways. Members of the IL-6, IL-11, and LIF signaling pathways were found to be enhanced during *S. aureus* infection in WT cells but remained downregulated in both infected and uninfected YAP^-/-^ cells. These cytokines support the proliferation and differentiation of hematopoietic stem cells ^41^, and LIF has been shown to enhance the killing of *S. aureus* by neutrophils ^42^. Although *IL6* was expressed at low levels in the nCounter panel, RT-qPCR showed that *IL6* expression was low in HEK293 cells but was nevertheless increased during *S. aureus* infection in WT cells and remained lower in both infected and uninfected YAP^-/-^ cells (**Figure 6C**). The expression of chemokine genes, including *CXCL8*, *CCL20*, *CXCL2*, and *CXCL1*, which are known to enhance immune cell recruitment and are consequently involved in *the S. aureus* inflammatory response, especially CXCL8, which is critical for neutrophil recruitment ^43, 44^, was enhanced during *S. aureus* infection in WT cells. Even if these genes were upregulated during *S. aureus* infection in YAP^-/-^ cells, the expression of these genes remained strongly downregulated in YAP^-/-^ cells compared to WT cells (**Figure 6AB**). Moreover, *S. aureus* infection enhanced the expression of *PTGS2* (also known as cyclooxygenase-2) (**Figure 6AB**), which encodes a key enzyme for the synthesis of prostaglandins that are involved in the inflammatory response against *S. aureus* ^45^; however, *PTGS2* expression remained lower in YAP^-/-^ infected and uninfected cells than in WT infected and uninfected cells (**Figure 6AB**). The inflammasome response is important during *S. aureus* infection for neutrophil recruitment ^46^. We found that several inflammasome-related genes, such as *CASP4* and *NLRC4*, were downregulated in YAP^-/-^ infected and uninfected cells. In addition, using RT-qPCR detect the low-level expression of *IL1B* was detected in WT cells during infection, whereas it remained undetectable in YAP^-/-^ infected and uninfected cells (**Figure 6C**). Although YAP/TEAD itself could contribute to the expression of cytokines and chemokines, we found that the expression of some transcription factors involved in inflammation was modified in YAP^-/-^ cells. During infection, nuclear factor κB (NF-κB) and activator protein 1 (AP-1) are known to trigger the first inflammatory response in cells ^47^. Our results confirmed that S. *aureus* infection triggers NF-κB pathway-related genes but does not increase NF-κB subunits (**S. Figure 6AB**). Likewise, we showed that *S. aureus* infection upregulates MAPK pathway-related genes with increased expression of AP1 members *JUNB* and *FOS* (**S. Figure 6AB**). However, in YAP^-/-^ control or infected cells, we found that these two pathways were highly disrupted due to the downregulation of genes encoding NF-κB subunits (*NFKB1, NFKB2, REL, RELA,* and *RELB*) and AP1 members, including *JUN, JUNB,* and *FOS* (**S. Figure 6A-B**). Of note, several other inflammatory pathways were altered in YAP^-/-^ cells, with a reduction in interferon signaling, NLR signaling, DNA sensing, and MHC class I signaling (**Figure 3A**). Altogether, these results highlight that YAP activity can modulate the expression of a wide range of inflammation-related genes involved in the response against *S. aureus*.

**Figure 6.**
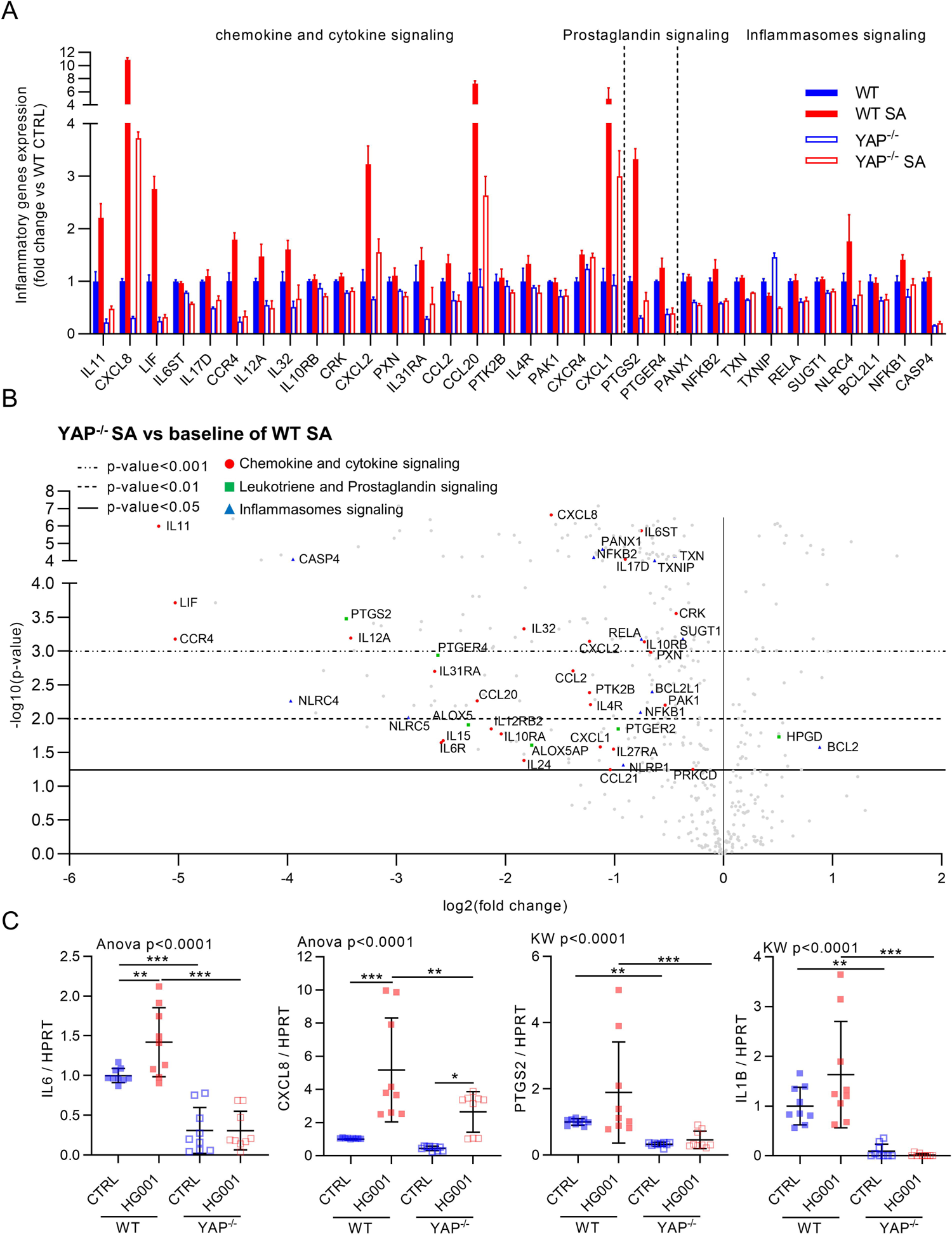
YAP promotes inflammatory response during *Staphylococcus aureus* infection. HEK293 cells were cultured and infected, as described in Figure 3. A: nCounter NanoString host response inflammatory gene expression in the four groups; depicted genes were selected if at least one comparison between two groups gave a corrected p-value<0.01, and they must be related to chemokine, cytokine, prostaglandin, or inflammasome signaling as indicated. B: Volcano plot representation of differential gene expression in YAP^-/-^ infected group versus the baseline of WT infected group; depicted genes are chemokine and cytokine (red circle), leukotriene and prostaglandin (green square), and inflammasome (blue triangle) pathway genes differentially expressed. C: RT-qPCR quantification of IL6, CXCL8, PTGS2, and IL1B expression normalized to HPRT expression. Results are expressed as fold change vs. control group (only in A and C) and presented as histograms (A) or individual values (C) with mean ± SD, representing three independent experiments (C). WT: Wild type; SA: *S. aureus*. Analysis of variance (ANOVA) test with false discovery rate (FDR) correction for multiple comparisons post hoc tests: * p<0.05, ** p<0.01, *** p<0.001.

### *S. aureus* ST80 infection in synovial organoids also modulates YAP signaling

Organoid-based infection models enable the study of infection in a 3D cell model using primary cells that display more physiological characteristics than continuous cell lines. In this experiment, we used a model of synovial organoids ^36^, formed with fibroblast-like synoviocytes (FLSs) from three different donors that were infected with ST80 strains. Given that *S. aureus* is one of the leading causes of osteoarticular infection in humans (Tong et al., 2015), this infection model should be highly clinically relevant.

Live-cell confocal microscopy showed that *S. aureus* was internalized in FLS and replicated in these cells (**Figure 7A**). We found that the EDIN-B-expressing ST80 WT strain thoroughly altered the organization of the actin cytoskeleton as early as 30 min post-infection and prevented the formation of actin fibers, whereas actin fibers were neither disrupted nor hindered by the ST80 Δ*edin*B strain (**Figure 7A**), which reflects the ability of EDIN-B to inhibit RhoA.

**Figure 7.**
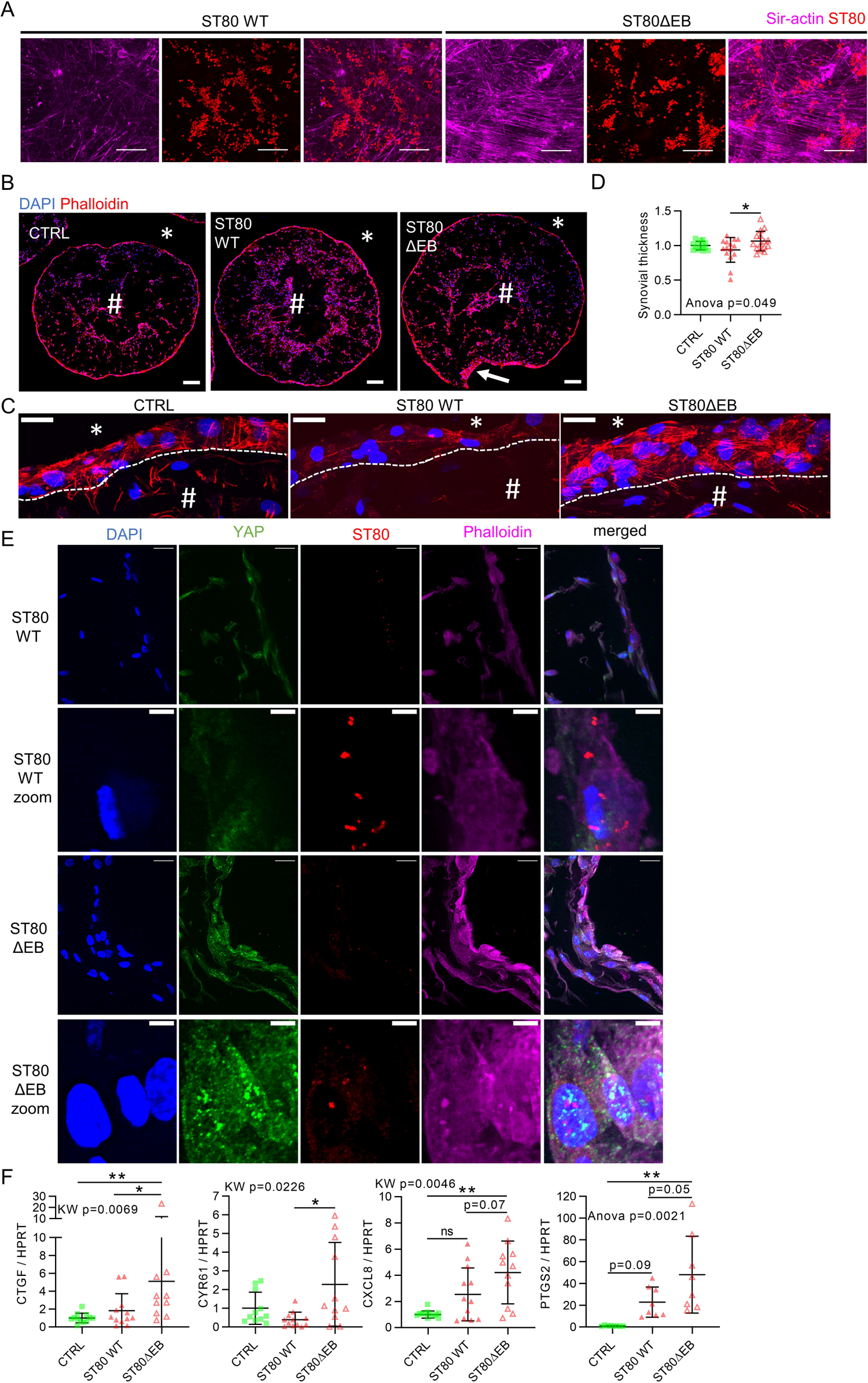
*Staphylococcus aureus* EDIN-B expression decreased YAP activity in a synovial organoid model. Primary fibroblast-like synoviocytes (FLSs) from human osteoarthritic patients (n=3) were used to form synovial organoids. Organoids were infected with 1×10^8^ *S. aureus* per mL. *Staphylococcus aureus* ST80 strains were allowed to contact organoids for 30 min and imaged immediately (A) or let in contact for 2 h upon agitation, then, lysostaphin was added at 10 µg/mL for the rest of the experiments to avoid extracellular *S. aureus* multiplication. A: Representative deconvolved confocal z-stack max intensity projection images of live organoids labeled with Sir-actin (actin filaments, magenta), ST80 strains (DsRed, red), and merged images were obtained from the surface of the organoid; scale bar: 25 µm. B-C: Confocal representative z-stack maximum intensity projection of 30 µm thick cryosections of the entire organoid (B) or the lining layer (C); #: organoid stromal part, *: outside of the organoid, dotted line: limit between lining layer and stromal part, white arrow: local synovial thickening; Scale bar: 200 µm (B), 25 µm (C). D: Microscopy quantification of synovial lining layer thickness. n=14 to 15 per group. E: Confocal representative z-stack max intensity projection images of 30 µm thick cryosections labeled with YAP (immunolabeling, green), DAPI (nucleus, blue), ST80 strains (DsRed, red), phalloidin (actin filaments, magenta), and merged; scale bar: 20 µm or 5 µm for zoom. F: RT-qPCR quantification of CTGF, CYR61, CXCL8, and PTGS2 expression normalized to HPRT expression, n= 11 to 12 per group. Results are expressed as fold change vs. control group and presented as individual values with mean ± SD, and representative of three independent experiments corresponding to three different fibroblast-like synoviocytes (FLS) donors. WT: wild type; ST80ΔEB: ST80 EDIN-B-deleted strain. Analysis of variance (ANOVA) or Kruskal-Wallis (KW) test with false discovery rate (FDR) correction for multiple comparisons post hoc tests: * p<0.05, ** p<0.01, *** p<0.001.

In this model of synovial organoids, FLS forms a lining layer at the edge of the organoid, and more sparse cells in the core of the structure are organized like a stroma mimicking the human synovial membrane ^36^ (**Figure 7B**). Synovial lining layer thickening, which is recognized as a hallmark of synovial inflammation, was found to be mediated through YAP activity ^36, 49, 50^. At 7 hpi, ST80 Δ*edin*B induced synovial lining layer thickening (**Figure 7B-D**) which suggest that YAP is activated. In contrast, ST80 WT that expressed EDIN-B did not induce an increase in of the synovial lining layer thickness (**Figure 7B-D**), probably reflecting YAP inhibition. This assumption was strengthened by the confocal microscopy images showing that the EDIN-B-expressing ST80 WT strain, unlike the ST80 Δ B strain, was able to reduce YAP immunolabeling in infected synovial lining layer cells at 7 hpi (**Figure 7E**). In addition, the ST80 WT strain was found to inhibit YAP transcriptional activity by reducing the expression of the connective tissue growth factor (*CTGF*) and *CYR61* genes, whereas the ST80 Δ*n*B strain increased their expression (**Figure 7F**), which confirmed that the EDIN-B-expressing ST80 WT strain was able to inhibit YAP transcriptional activity in synovial organoids. In addition, since the *CXCL8* and *PTGS2* genes were downregulated in HEK293 YAP^-/-^ infected cells compared to WT infected cells, we assessed the expression of these two genes in synovial organoids. Compared to uninfected organoids, ST80 Δinduced an increase in the expression of *CXCL8* and *PTGS2*, in contrast to the EDIN-B-expressing ST80 WT strain, which was consistent with YAP transcriptional activity inhibition in HEK293 cells.

In conclusion, this organoid-based model confirmed the strong ability of the EDIN-B-expressing ST80 WT strain to inhibit YAP transcriptional activity and reduce the expression of inflammatory mediators. These results suggest that YAP inhibition by EDIN-B can reduce synovial inflammation and prevent immune cell recruitment at the infection site.

## DISCUSSION

In this study, we sought to investigate whether YAP is involved in the clearance of intracellular *S. aureus*. We demonstrated that YAP plays a critical role in efficient cell-autonomous immune response against intracellular *S. aureus* by controlling autophagy-lysosome and inflammation-related signaling pathways.

Our results are consistent with the role of YAP/TAZ transcriptional activity in promoting autophagy ^14, 15, 51^. In our model of YAP-mutated cells, we found no differences in *MLC2* and *DIAPH1* expression, which is important for autophagosome formation ^14^. In contrast, we observed alterations in the late phase of autophagy, which is consistent with a previous study showing impaired fusion between autophagosomes and lysosomes ^15^, but the mechanism involved seems to be different. Indeed, we observed abnormal and oversized autophagic vesicles. In particular, we found that YAP^-/-^ cells have an alteration in lysosomal acidification, which can be explained by the decreased expression of *ATP6V0D1*, which encodes a V-ATPase subunit required for lysosomal acidification ^52^. Moreover, abnormal and oversized autophagolysosomes with poor degradative functionality have been described in cells deficient for V-ATPase subunits ^39^, which supports our findings. In addition, we found that *CTSL* was downregulated in the YAP^-/-^ HEK293 cells. It has been shown that *CTSL*-deleted cells have important lysosomal dysfunction and LC3-II accumulation, reflecting an altered autophagic flux ^38^. In addition, we showed that this effect was mediated by the YAPΔTEAD binding domain, since YAP TEAD^-/+^ cells display autophagic defects similar to those of YAP^-/-^ cells. However, we cannot exclude the possibility that YAP interacts through its TEAD-binding domain with another transcription factor involved in autophagy and lysosome signaling pathways. Indeed, a recent study showed that YAP can interact with transcription factor EB (TFEB) to induce the expression of autophagic and lysosomal genes ^51^ but whether the TEAD-binding domain of YAP is required for its interaction with TFEB is unknown. Thus, YAP/TEAD and/or YAP/TFEB could act synergistically for autophagy-and lysosome-related gene regulation. Overall, our data reinforce the role of YAP in autophagy regulation and provide new insights into how YAP promotes autophagic flux.

Our work shows that YAP transcriptional activity is required to control the replication of *S. aureus*, and that some EDIN-B-expressing *S. aureus* strains can inhibit YAP to promote their own intracellular replication. Given that autophagy is clearly established as a major mechanism for clearing *S. aureus in vitro* and *in vivo* ^31–33^, the autophagy dysfunction observed in YAP-mutated cells in this study could explain why *S. aureus* infection was more pronounced when YAP transcriptional activity was inhibited or absent. The altered lysosomal function observed in YAP-mutated cells may also decrease *S. aureus* clearance as *CTSL*-deficient macrophages exhibit a poor ability to remove intracellular *S. aureus* ^53^, and genetic manipulations of V-ATPases or bafilomycinA1 treatment in macrophages promoted *S. aureus* intracellular replication ^54^. However, to avoid degradation, *S. aureus* has also been found to inhibit the fusion of autophagosomes with lysosomes and escape from autophagic vesicles to replicate inside the cytosol ^31^. In our model, the loss of YAP transcriptional activity, which induces the blockage of autophagic flux, was found to promote autophagosome escape and replication of *S. aureus*.

In addition, we found that EDIN-B-expressing *S. aureus* inhibited YAP transcriptional activity, which enabled them to replicate more efficiently in the cells, likely by escaping from autophagic vesicles. Autophagy is a conserved cellular process known to be involved in the clearance of intracellular bacteria ^55^, and RhoA-targeting toxins (such as EDINs) can be expressed by other pathogenic bacteria such as *Yersinia* and *Salmonella* species ^24, 25^. In addition, bacteria secreting RhoA targeting-toxins were found to alter actin dynamics, leading to the impairment of tight and adherent junctions and an increase in bacterial invasion across the epithelium and endothelium ^25, 27^. Interestingly, YAP is known to promote the formation of focal adhesion complex, and regulate actin dynamics, and be activated after intestinal barrier disruption following bacterial infection ^14, 21, 56^. Thus, we speculated that some known RhoA inhibition mechanisms achieved by bacteria could be mediated by YAP activity.

Increasing evidence demonstrates that YAP/TEAD transcriptional activity can play a pro-inflammatory role by promoting the expression of pro-inflammatory mediators such as *IL6* ^12^, *CCL2* ^57, 58^, *IL8* ^9^, *IL1B* ^13^, *PTGS2* ^59^, and NF-kB family members ^60^. However, contradictory results exist in the literature, indicating an anti-inflammatory role of YAP in mouse models ^20, 61, 62^. In our study, YAP transcriptional activity was found to have a pro-inflammatory effect in HEK293 cells. We found that the loss of YAP activity decreased the expression of several pro-inflammatory genes known to foster *S. aureus* clearance *in vivo*. Thus, RhoA-mediated inhibition of YAP by bacteria could be a way to evade the immune system by decreasing the inflammatory response.

Another important question is how bacteria modulate YAP activity in host cells. In contrast to our results, *S. aureus* infection in *Drosophila* was found to increase Yorkie cytoplasmic localization in fly fat bodies ^22^. Yorkie overexpression in fly fat bodies was found to increase *S. aureus-*induced death compared to WT flies ^22^. However, there are important differences between human NPPCs and fly fat bodies, which can influence YAP activity and its subcellular localization upon infection. In mice, several other bacterial species (*e.g.*, *Streptococcus pneumoniae* and *Helicobacter pylori*) lead to nuclear translocation ^18, 20^. Thus, it could be interesting to test whether YAP has an anti-*S. aureus* function in mouse models.

Although bacteria-induced tissue damage can promote YAP activation in mouse models, the mechanisms that induce nuclear translocation of YAP upon bacterial infection are not fully understood ^20, 21^. Our results showed that *S. aureus* supernatant alone is not sufficient to induce YAP nuclear translocation, which suggests that internalization of *S. aureus* is needed for inducing nuclear translocation of YAP. Interestingly, *S. aureus* internalization is mainly driven by α 1 integrins, which trigger the activation of focal adhesion kinase (FAK). YAP is known to be highly sensitive to cell mechanical stimulation, such as integrin-FAK activation, which increases RhoA activity and causes YAP nuclear translocation ^63^. Thus, it will be important to investigate whether YAP activation following *S. aureus* internalization could be a nonspecific “danger signal” by converting cell mechanical events into cell-autonomous immune responses, including xenophagy and inflammatory responses.

Overall, this work provides new fundamental insights into the role of YAP in cell-autonomous immune responses. It also provides new insight into the role of the C3 exoenzyme EDIN during *S. aureus* infections. Thus, the findings of this work could help find new ways to fight intracellular bacteria and open the way for future microbiology and YAP-related investigations.

## Methods

### Cell culture

HEK293 cells were cultured in Dulbecco’s modified Eagle’s medium (DMEM, Sigma-Aldrich, St. Louis, MO, USA) with 10% fetal bovine serum (FBS), 1% non-essential amino acid solution, and 1% penicillin and streptomycin (PS) solution. The plates were coated with fibronectin (1:100, Sigma-Aldrich, F1141) for 2 h at 37 °C before use. HEK293 cells were grown at different cell densities: For low density (LD) cell culture, cells were seeded at 10,000 cells/cm^2^ and used 24 h after seeding; for medium density (MD) cell culture, cells were seeded at 100,000 cells/cm^2^ and used 24 h after seeding; for high density (HD) cell culture, cells were seeded at 100,000 cells/cm^2^ and used 72 h after seeding.

### Cell line generation using CRISPR-Cas9 technology

HEK293 YAP^-/-^ were generated using commercially available plasmids with specific CRISPR-Cas-9 single guide RNA (sgRNA) and sequence for homology-directed repair targeting YAP sequence (Santa Cruz Biotechnology, Dallas, TX, USA) as previously reported ^36^. HEK293 YAPΔTEAD^-/+^ cells were generated using the CRISPR-Cas9 technique and homology-directed repair. sgRNA was designed to cut in exon 1 of the YAP gene at proline 98 using the following protospacer: 5′ CGACTCCTTCTTCAAGCCGC-3′ Homologous recombination was supported by a donor plasmid with a 5′ homology arm of 681 bp, a 3′ homology arm of 837+12 bp, whose original sequence TTCAAGCCGCCG was modified by the sequence AGAAGAAGAAGA that introduced the following mutations: Phe96Arg, Lys97Arg, Pro98Arg, and Pro99Arg. CRISPR-Cas9 and donor plasmids were manufactured on demand by VectorBuilder (VectorBuilder, Neu-Isenburg, Germany). HEK293 cells were transfected with 0.5 µg of each plasmid and 2 µL transfection reagent (Jet prime, Polyplus transfection, New York, NY, USA) in a final volume of 100 µL. After 48 h of transfection, the cells were seeded at one cell per well in a 96-well plate for monoclonal expansion. Mutations following homologous recombination were confirmed by PCR sequencing (Eurofins Genomics, Nantes, France).

### Bacterial strains and plasmids

*Staphylococcus aureus* strains used in the study were the HG001 strain, which is a methicillin-susceptible *S. aureus* (MSSA) strain that lacks edin genes ^64^ and the LUG1799 strain, which is a minimally passaged strain belonging to the European lineage community-acquired methicillin-resistant *S. aureus* (CA-MRSA) ST80-MRSA-IV strain ^65^ and is referred to as ST80 wild-type (WT) and its isogenic *edin*B mutant that is referred to as ST80 Δ^27^. All strains were stored at −20°C in cryotubes.

For live cell imaging, the plasmid pSK265, a derivative of pC194 ^66^, was used to express the *DsRed* gene under the control of the *rpob* promoter in *S. aureus* strains. All strains were transformed with the plasmid pSK265::DsRed by electroporation (Gene Pulser, Bio-Rad) and were grown at 37 °C on blood agar (43049, Biomérieux) or tryptic soy agar (TSA) (920241, Becton Dickinson) supplemented with 20 µg/mL of chloramphenicol when appropriate.

### Organoid culture and processing

Synovial organoids were assembled as previously described ^67^ with modifications ^36^. Fibroblast-like synoviocytes (FLS) were collected from osteoarthritis (OA) patients who provided written consent after oral information (IRB # 2014-A01688-39). FLS were mixed in phenol red-free Matrigel (356237, Corning, Corning, NY, USA) at 4×10^6^ cells/mL, and a single 22 µL droplet (representing approximately 90,000 cells) was added to each well of a 96-well U-shaped very low-attachment surface plate (CLS4515, Corning). The plate was incubated at 37 °C in 5% CO_2_ for 45 min to allow droplet gelation. Wells containing solidified droplets were filled with 200 µL of DMEM high-glucose medium supplemented with 10% FBS, 1 % glutamine, 1 % nonessential amino acids, 1 % PS, 0.1 mM ascorbic acid, and insulin (10 µg/mL)-transferrin (10 µg/mL)-selenium (3×10^-8^ M) solution at 37 °C in 5% CO_2_ for 21 days. At day 21, organoids were fixed with glyoxal solution at pH 4.5 (e.g., for 500 mL: 355 mL ddH_2_O, 99 mL ethanol, 39 mL glyoxal (128465, Sigma-Aldrich), and 1 mL acetic acid) for 1 h at room temperature (RT) because PFA fixation was deleterious. Organoids were embedded in a gelatin 100G (7.5%)-sucrose (10%) solution and frozen in an isopentane bath at −50 °C for 2 min before storage at −80 °C.

### Bacterial infection of HEK293 cells and organoids

HEK293 cells and organoids were infected with *S. aureus* using the enzyme protection assay (EPA) technique as previously described ^34^. Briefly, *S. aureus* bacterial suspensions were adjusted to an OD_600_ of 0.5 and serially diluted in the culture media of HEK293 cells or organoids. HEK293 cells were infected at a multiplicity of infection (MOI) of 1 (or 10 if indicated) for 2 h at 37 °C and 5% CO_2_. Organoids were infected with 1×10^8^ *S. aureus* per well in 24-well plates for 2 h at 37 °C and 5% CO_2_ with gentle agitation. After incubation, media was replaced with fresh culture media supplemented with 10 µg/mL lysostaphin (Ambicin, Ambi Products, Lawrence, NY, USA) to kill extracellular *S. aureus*. Bacterial suspensions used for infection challenges were seeded on agar plates and quantified after a 24-h incubation period to verify the real bacterial concentration. To quantify the intracellular load of *S. aureus* by culture, HEK293 cells were washed with phosphate buffered saline (PBS) to remove lysostaphin. Cells were lysed by osmotic shock using lysis buffer containing 0.25% Triton X-100 (Sigma-Aldrich), 0.25X trypsin-EDTA (Sigma-Aldrich), and sterile water. The *S. aureus* load of cell lysates was quantified on an agar plate using an automatic plate seeder (EasySpiral Dilute, Interscience, St-Nom la Bretèche, France) and a colony counter (Scan 4000, Interscience).

### Immunofluorescence

HEK293 cells were fixed with 4% PFA at RT for 20 min or in ice-cold methanol for 15 min (for LC3A/B immunolabeling). Fixed and frozen organoids were cryosectioned to a thickness of 30 µm. Samples (cells or cryosections) were rehydrated in PBS for 10 min and permeabilized in 0.3% Triton X-100 for 15 min. The samples were then incubated in blocking buffer containing 1% BSA, 5% goat serum, and 0.1% Triton-X100 for 60 min at RT. Subsequently, the samples were incubated with the primary antibody or isotypic control diluted in blocking buffer overnight at 4 °C. The antibodies used were mouse IgG anti-YAP antibody (63.7 sc-101199, Santa Cruz Biotechnology; 1:100), rabbit anti-LC3A/B antibody (4108, Cell Signaling Technology, Leiden, The Netherlands), mouse and rabbit IgG isotype antibody (31903 and 31235, Thermo Fisher Scientific; used at the same concentration as YAP or LC3A/B antibodies). After washing, the cells were incubated with secondary antibody, goat anti-mouse 488 or goat anti-rabbit 488 diluted in blocking buffer (A11034 and A32731, Thermo Fisher; 1:400) for 75 min at RT. The cells were counterstained with 4′,6-diamidino-2-phenylindole (DAPI) for 10 min at 37 °C with or without dye-labeled phalloidin (ab176753 or ab176759, Abcam, Cambridge, UK) for 1 h at 37 °C. For LC3 immunolabeling (ref, Cell Signaling Technology, Leiden, The Netherlands), the cells were fixed with ice-cold methanol for 15 min, and the immunolabeling procedure was identical to YAP immunolabeling.

### Live-cell confocal microscopy of HEK293 cells or organoids

Cells and organoids were infected with DsRed-expressing *S. aureus* strains using the enzyme protection assay (EPA) technique described above. In HEK293 cells, autophagosomes were labelled using the CYTO-ID Autophagy Detection Kit 2.0 (ENZ-KIT175, Enzo Life Sciences) as recommended by the manufacturer. Briefly, 30 min before image recording (*i.e.*, 2.5 hpi or 6.5 hpi), the spent media was discarded, and cells were washed once with the assay buffer. Cells were incubated with the CYTO-ID Green detection reagent and 5 µg/mL Hoechst 33342 for 30 min at 37 °C and 5% CO_2_ protected from light. Cells were then washed once with the assay buffer and imaged immediately by confocal microscopy. In organoids, Actin-F was labelled with Sir-Actin dye (1:5000, Cytoskeleton, Denver, CO, USA) for 4 h prior to infection.

### Image acquisition and quantification

Images were acquired using a spinning disk confocal microscope (SDCM) (Ti2 CSU-W1, Nikon, France) with a 60x objective (CFI Plan Apo Lambda NA = 1.40, MRD1605, Nikon) or using a confocal laser scanning microscope (CLSM) (LSM 800 airyscan, Zeiss, Oberkochen, Germany) with a 10x objective (Plan-Apochromat 10x/0.45 M27, Zeiss). Image analysis was performed with the General Analysis 3 module of the NIS software (v5.30, Nikon) or Fiji software (v1.52p, NIH, USA).

In HEK293 cells, YAP immunolabeling was quantified using an automatic macro developed with the NIS software to measure MFI in the cytoplasmic and nuclear areas and to calculate the NC ratio by dividing the nuclear MFI by the cytoplasmic MFI. CytoID quantification was also performed using the NIS software. Briefly, the images were denoised and binarized in 3D. The CytoID vesicle count and volume as well as the *S. aureus* volume were measured, and the colocalization between the *S. aureus* volume and CytoID vesicle volume was assessed. The cell area was determined using an extended area of DAPI labeling. For each cell, the count and mean volume of the CytoID vesicles were measured. The same method was used to measure the *S. aureus* volume per cell. Quantifications were performed by analyzing 2 to 3 fields per well using a 60x objective.

For organoid lining layer thickness, quantification was performed with the Fiji software using cryosections stained with DAPI and dye-labeled phalloidin. Two slices per organoid were assessed. Quantification was performed on tile images acquired with a 10x objective, allowing quantification of the entire structure. Images were binarized, and the synovial lining layer area was automatically selected. The organoid perimeter was then measured. The lining layer thickness was the result of the synovial lining layer area divided by the perimeter of the synovial organoid.

### Luciferase assay

HEK293 cells were transfected in 96-well plates with the 8xGTIIC-luciferase plasmid (firefly luciferase, # 34615, Addgene, Watertown, MA, US) and the pRL-SVl40P plasmid (Renilla luciferase, # 27163, Addgene), using 0.5 µg of each plasmid and 2 µL of the jetPRIME transfection reagent (Polyplus transfection, New York, NY, USA) in a final volume of 100 µL per well and incubated overnight at 37°C in 5% CO_2_. The next day, the spent medium was replaced with the fresh complete culture medium, and the cells were incubated for another 24 h at 37°C in 5% CO_2._ The day after, the transfected cells were challenged with *S. aureus* or supernatant only, as mentioned in the text. After the challenge, the cells were lysed and luminescence was quantified using the Promega dual glow assay (Promega, Madison, WI, USA) with a multimodal plate reader (TriStar, Berthold). The blank value was subtracted, and the firefly luciferase activity was divided by the Renilla luciferase activity to normalize the results according to the number of cells.

### Protein extraction and western blotting

For HEK293 cells, protein extraction was performed using the Allprep RNA/Protein Kit (80404 Qiagen Inc., Hilden, Germany). Proteins (10–20 µg) were denatured and separated for 20 min at 200 V before being transferred onto the polyvinylidene difluoride membrane (IB24002, Thermo Fisher Scientific). The membrane was blocked in TBS Tween 0.1% with 5% skimmed milk and incubated with primary antibody overnight at 4 °C. The membrane was washed once with washing buffer and incubated with a horseradish peroxidase-conjugated secondary antibody (31460, Thermo Fisher Scientific; 1:5000) for 1 h at RT. Immunoreactive protein bands were visualized using the Clarity Western ECL Substrate (Bio-Rad, Hercules, CA, USA). Western blotting (WB) was performed using the following primary antibodies purchased from Cell Signaling Technology (Danvers, MA, USA) diluted at 1:1,000: YAP/TAZ (#8418), LC3A/B (#12741), and 1:5,000: GAPDH (#2118).

### RNA extraction and RT-qPCR

For synovial organoids, lysis was performed using the TRI Reagent (Sigma-Aldrich); three synovial organoids were pooled together during the lysis step to yield sufficient RNA. For synovial organoids, the aqueous phase was processed following lysis in the TRI Reagent for RNA extraction and purification. For cell culture, RNA was extracted using the Allprep RNA/Protein Kit (Qiagen). The quality and quantity of RNA were assessed using the Experion RNA Analysis Kit (BioRad) and QuantIT RiboGreen RNA Assay Kit (Thermo Fisher Scientific), respectively. Complementary DNA (cDNA) was synthesized using an iscript cDNA Synthesis Kit (Bio-Rad). Quantitative RT polymerase chain reaction (PCR) was performed using the CFX96 RealTime System (BioRad) with LightCycler FastStart DNA Master plus SYBR Green I (Roche Diagnostics, Basel, Switzerland). The results were normalized to the housekeeping gene expression hypoxanthine-guanine phosphoribosyltransferase (HPRT). The sequences of the primers used in this study are available upon request.

### Transcriptomic analysis using nCounter Host Response Panel

The nCounter Host Response panel (Nanostring technology), which includes 770 genes involved in host response processes, was performed with the nCounter Sprint instrument following the manufacturer’s recommendations. Briefly, we used 50 ng of RNA extracted from WT or YAP^-/-^ HEK293 infected (or not) with the HG001 strain at MOI 10 for 7 h (n = 3 per group). All quality controls were performed according to the manufacturer’s instructions. Normalization was performed using the housekeeping genes identified by the geNorm analysis using the NanoString advance software. The count detection limit was determined using a threshold based on the negative controls. Data analysis was performed using the nSolver package (version 3.0) and the Advanced Analysis module (version 1.0.36). Differential expression and pathway analyses were performed using the nSolver advance analysis module according to the guidance given by manufacturer’s instructions. Genes with a false discovery rate (FDR)-corrected p-value < 0.05 were considered significantly differentially expressed.

## Statistical analysis

Data are represented as single values with mean and standard deviation and are expressed, if indicated in the figure legend, as a percentage of the mean of control values. The results are representative of at least three independent experiments. Multiple comparisons were performed by analysis of variance (ANOVA) or Kruskal-Wallis test, and post hoc comparisons were corrected using the FDR method of Benjamini and Hochberg. Results were considered significantly different when p<0.05 or q<0.05. All statistical analyses were performed using the GraphPad software (v9.2.0, Prism). The NanoString results were analyzed using the nSolver software (v4.0, NanoString Technology) and nSolver Advance Analysis Module (v2.0.134, NanoString Technology).

## ACKNOWLEDGMENT

The strain of *Staphylococcus aureus* HG001 was provided by Tarek Msadek (Institu Pasteur, France). We thank all the members of the GIMAP and LBTO teams for their help and feedback.

## AUTHORS CONTRIBUTIONS

RC, EA, and POV designed the experiments. RC performed the experiments, designed and performed the CRISPR-cas9 based HEK293 mutation and developed the synovial organoid model. EA contributed to the plasmid design for YAPΔTEAD^-/+^ cell generation. MT contributed to the nanostring experiments and CRISPR clone identification. EA, YD, and KR contributed to protein extraction and WB experiments. AP-transformed *S. aureus* strains for live-cell microscopy. ED contributed to RNA extraction in the organoid experiments. FV provided ST80 WT and Δ*edin*B strains. RC and EA analyzed the results. RC, EA, and POV wrote the manuscript. JJ, HM, FV, and FL provided critical corrections to the manuscript. POV, FL, and JJ obtained funding for this work and supervised the project. All authors have agreed to the final version of the manuscript.

## DATA AVAILABILITY

The datasets generated and analyzed during the current study are available from the corresponding author upon request. Nanostring nCounter data have been deposit on Gene Expression Omnibus platform under the accession number: GSE197181.

