## Supplementary Figures for "YAP promotes cell-autonomous immune responses to tackle intracellular *Staphylococcus aureus in vitro*"

**SUPPLEMENTAL INFORMATION**

**LEGENDS OF SUPPLEMENTAL FIGURES**

**S. Figure 1. *S. aureus* HG001 strain do not inhibit YAP activity if YAP is already activated**

HEK293 cells were cultured at low density (A) or medium density (B-G). HG001 *S. aureus* infection was at a multiplicity of infection of 10 for 7 h (or 3 h if indicated). *S. aureus* were allowed to contact for 2 h with the cells, and lysostaphin was added at 10 µg/ml for the rest of the experiments to avoid extracellular *S. aureus* multiplication. A: Confocal representative z-stack max intensity projection images of YAP (immunolabeling, green), DAPI (nucleus, blue), *S. aureus* (DsRed, red), and phalloidin (actin filament, magenta). Scale bar: 5 µm. B-D: Representative western blot results of YAP, TAZ, and GAPDH (B) with their quantification normalized by GAPDH expression (C and D). E-F: Luciferase reporter assay of TEAD transcription factor activity (8xGTIIC) for HG001 *S. aureus* infection at 3 hpi (E) or 7 hpi (F). G: RT-qPCR quantification of CYR61 expression normalized to HPRT expression. Results were expressed as fold change vs. control group and presented as individual values with mean ± SD, representing three independent experiments. ANOVA or Kruskal-Wallis (KW) test with FDR correction for multiple comparisons post hoc tests: * p<0.05; *** p<0.001.

**S. Figure 2. *S. aureus* supernatant containing EDIN-B prevented YAP activation in HEK293 cells**

HEK293 cells were cultured at high density. The ST80 strain supernatant was added for 24 h. A: Confocal representative z-stack max intensity projection images of YAP (immunolabeling, green), DAPI (nucleus, blue), and merged. Scale bar: 20 µm. B-D: Quantification of YAP nuclear mean fluorescence intensity (MFI) (B), YAP cytoplasmic MFI (C), and YAP nuclear cytoplasmic ratio (D). Results were expressed as fold change vs. control group and presented as individual values with mean ± SD, representing three independent experiments. CTRL: control; WT: wild-type; ST80ΔEB: ST80 EDIN-B-deleted strain; Sp: supernatant. ANOVA test with FDR correction for multiple comparison post-hoc tests: *** p<0.001.

**S. Figure 3. Autophagy / lysosome pathways regulation during infection in WT and YAP^-/-^ cells**

HEK293 cells were cultured at medium density and infected with HG001 *S. aureus* strain at a multiplicity of infection of 10 for 7 h. *S. aureus* were let in contact for 2 h with the cells, then, lysostaphin was added at 10 µg/ml for the rest of the experiments to avoid extracellular *S. aureus* multiplication. A: nCounter Nanostring host response autophagy and lysosome gene expression in the four groups; depicted genes were selected if at least one comparison between two groups gave a corrected p-value<0.01. B: Volcano plot representation of differential gene expression in YAP^-/-^ infected group versus the baseline of WT infected group; depicted genes are lysosome (green) and autophagy (red) related pathway genes differentially expressed. C: RT-qPCR quantification of CTSL expression normalized to HPRT expression. WT: Wild type; SA: *S. aureus*. Results are expressed as fold change vs. control group and presented as histograms (A) or individual values (C) with mean ± SD, representing three independent experiments (C). ANOVA or Kruskal-Wallis (KW) test with FDR correction for multiple comparisons post hoc tests (C): * p<0.05; *** p<0.001. The p-value was calculated using nanostring advanced software based on t-test corrected with false discovery rate (B).

**S. Figure 4. YAP/TEAD transcriptional activity is required to reduce *S. aureus* induced autophagic flux blockage**

HEK293 cells were cultured at medium density. HG001 *S. aureus* infection was at a multiplicity of infection of 1 for 3 or 7 h, as indicated. *S. aureus* were allowed to contact for 2 h with the cells, and lysostaphin was added at 10 µg/ml for the rest of the experiments to avoid extracellular *S. aureus* multiplication. Representative confocal (0.5 µm thick z-stack) images of live cells labelled with DAPI (nucleus, blue), CytoID (LC3-II vesicles, green), HG001 (DsRed, red), and merged; scale bar: 5 µm. Note that WT infected cells at 7 hpi are also depicted and represent another infection scenario compared to Figure 5A. B-C: corresponding quantification of the LC3-II positive vesicle count (B) or mean volume (C) per cell as indicated. Each point represents one cell. The number of analyzed cells per group is shown. The results are expressed as individual values with mean ± SD, representing three independent experiments. Kruskal-Wallis (KW) test with FDR correction for multiple comparisons post hoc tests: * p<0.05, ** p<0.01, *** p<0.001.

**S. Figure 5. EDIN B expression by *S. aureus* promotes its escape from autophagic vesicles**

HEK293 cells were cultured at medium density. ST80 *S. aureus* infection was at a multiplicity of infection of 1 for 3 or 7 h, as indicated. *S. aureus* were allowed to contact for 2 h with the cells, and lysostaphin was added at 10 µg/ml for the rest of the experiments to avoid extracellular *S. aureus* multiplication. A: Representative confocal (0.5 µm thick z-stack) images of live cells labelled with DAPI (nucleus, blue), CytoID (LC3-II vesicles, green), HG001 (DsRed, red), and merged; scale bar: 5 µm. B: Quantification of the relative percentage of colocalization between *S. aureus* and CytoID labelling (LC3-II vesicles). The results were expressed as fold change vs. the WT 3 hpi group set at 100%. C-D: corresponding quantification of the LC3-II positive vesicle count (C) or mean volume (D) per cell as indicated. Each point represents one cell. The number of analyzed cells per group is shown. The results are expressed as individual values with mean ± SD, representing three independent experiments. T-test (B) or Kruskal-Wallis (KW) test with FDR correction for multiple comparisons post hoc tests: * p<0.05, ** p<0.01, *** p<0.001.

**S. Figure 6. YAP promotes inflammatory response during *S. aureus* infection**

HEK293 cells were cultured and infected, as described in Supplementary Figure 3. A: nCounter Nanostring host response for NF-κB and MAPK gene expression in the four groups; depicted genes were selected if at least one comparison between two groups gave a corrected p-value<0.01. Results are expressed as histograms with mean ± SD. B: Volcano plot representation of differential gene expression in YAP^-/-^ infected group versus the baseline of WT infected group; depicted genes are NF-κB (red circle) and MAPK (blue triangle) related pathway genes differentially expressed. The p-value was calculated using nanostring advanced software based on a t-test corrected with false discovery rate. WT: Wild type; SA: *S. aureus*.

**SUPPLEMENTARY FIGURES**

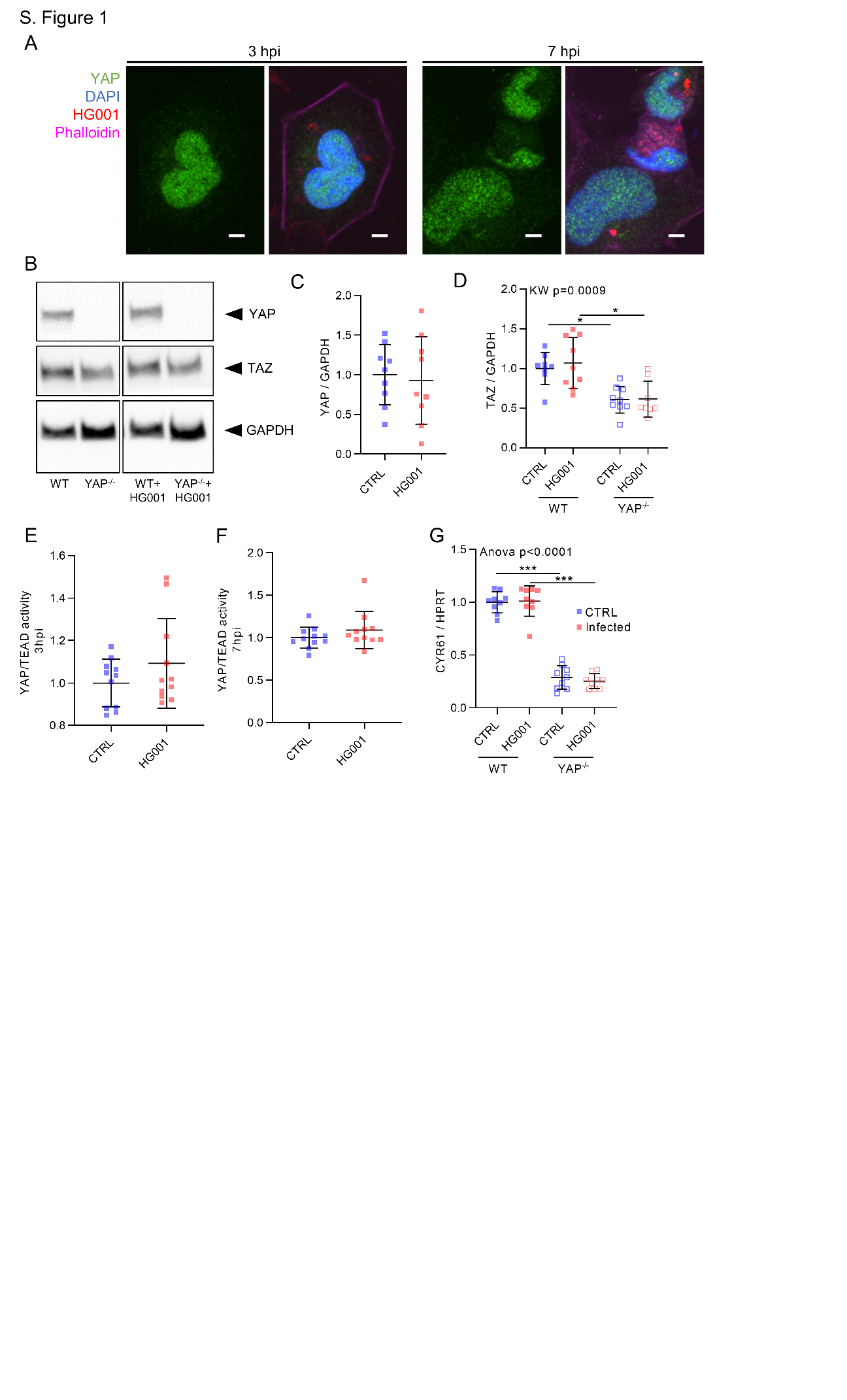

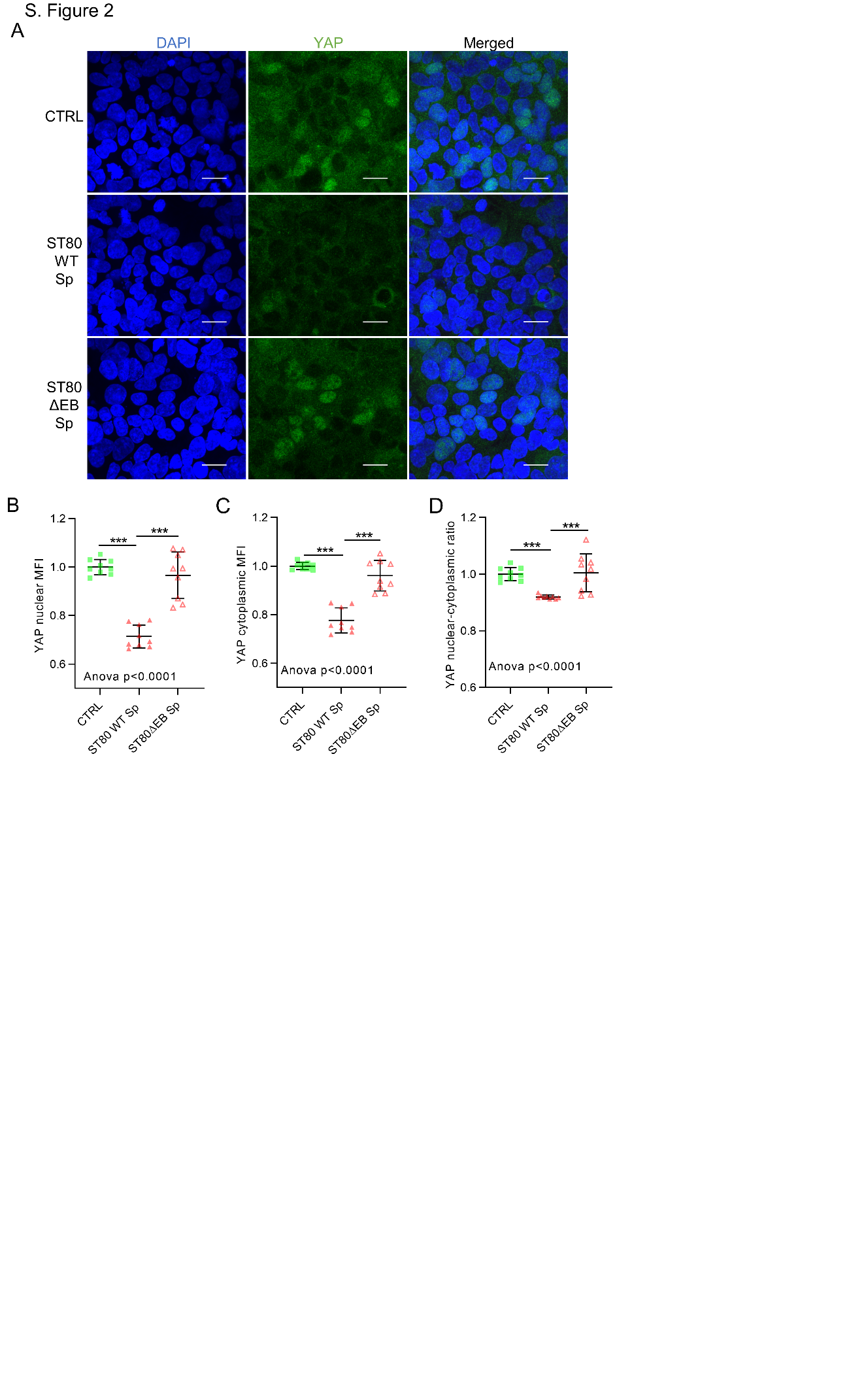

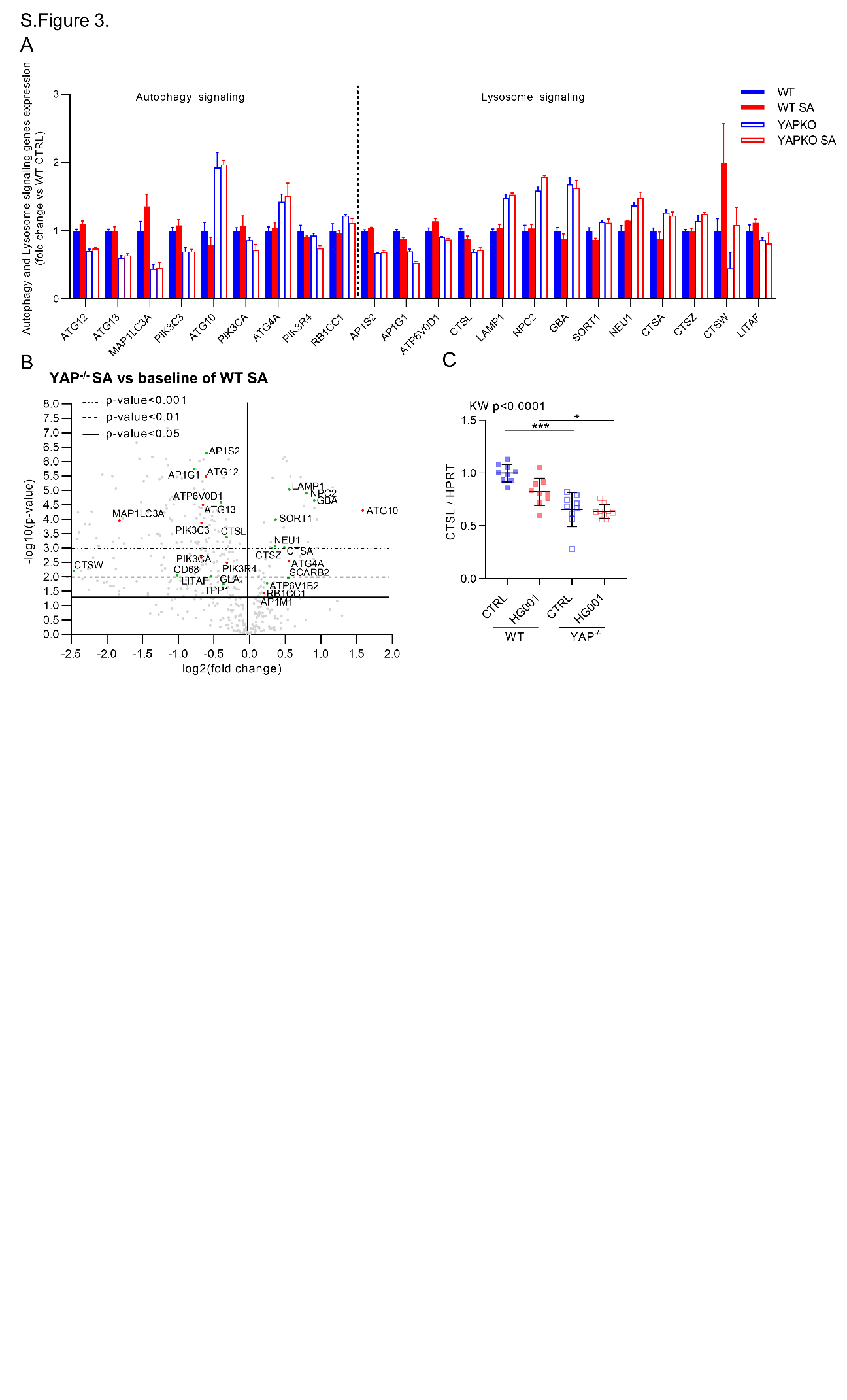

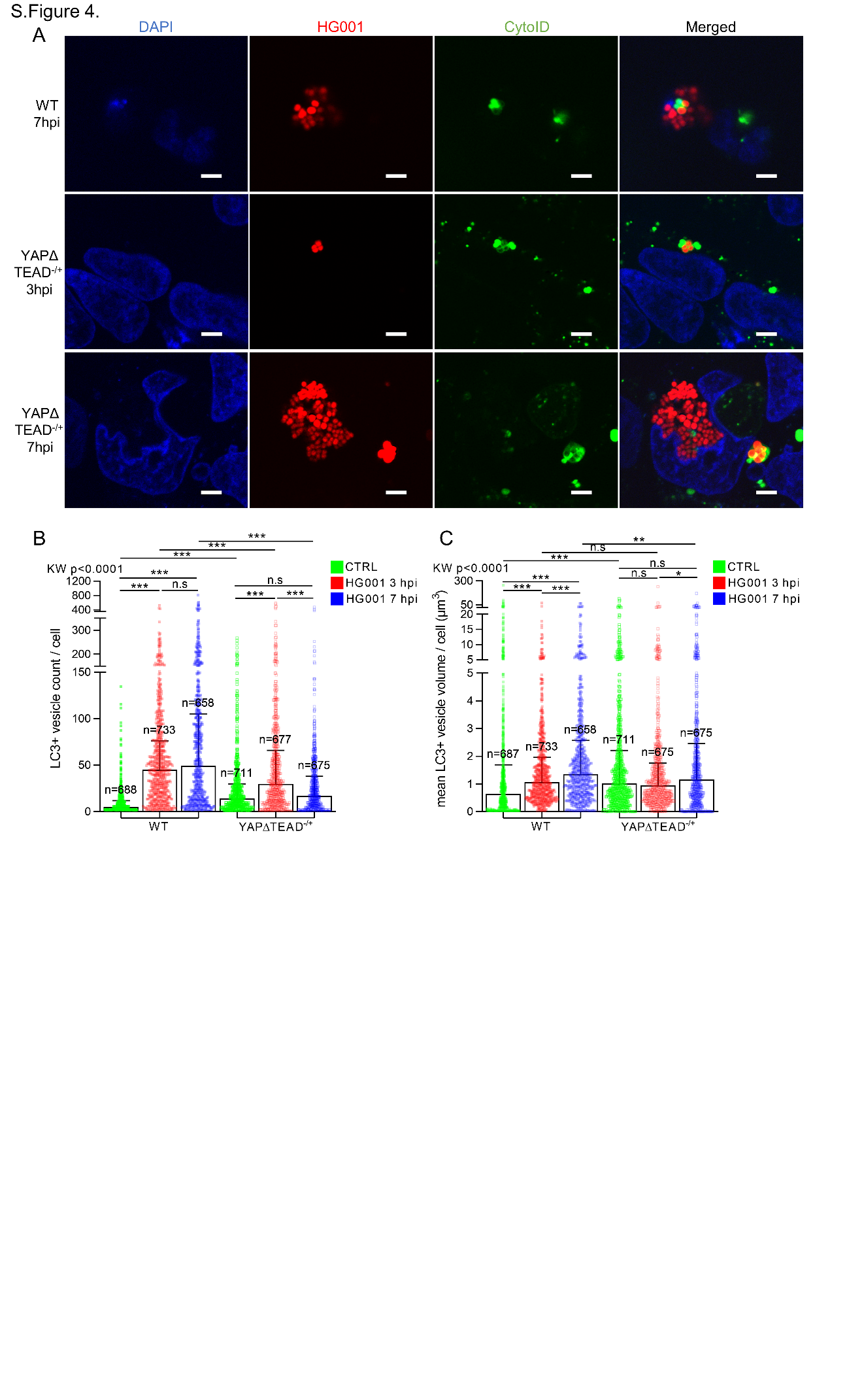

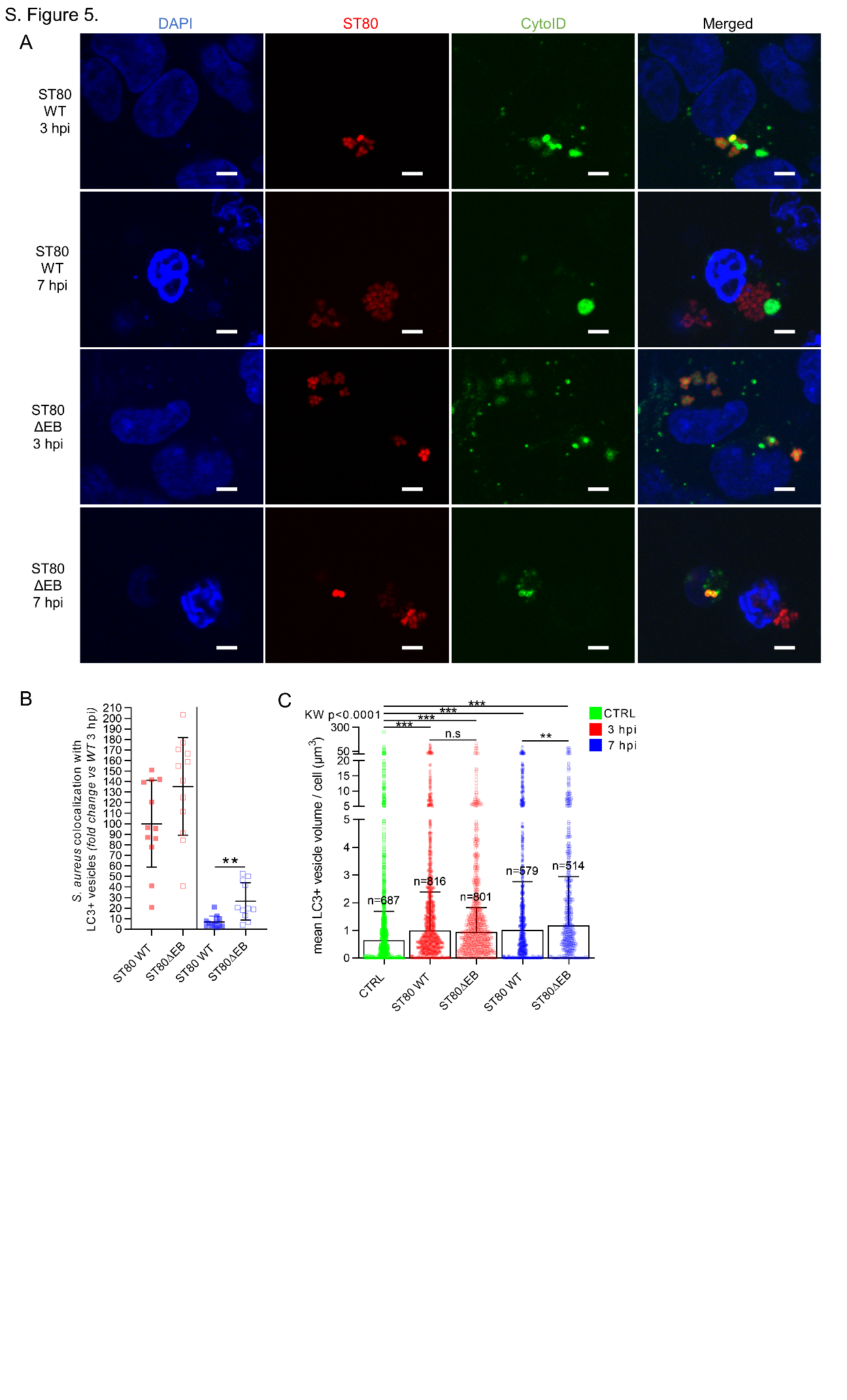

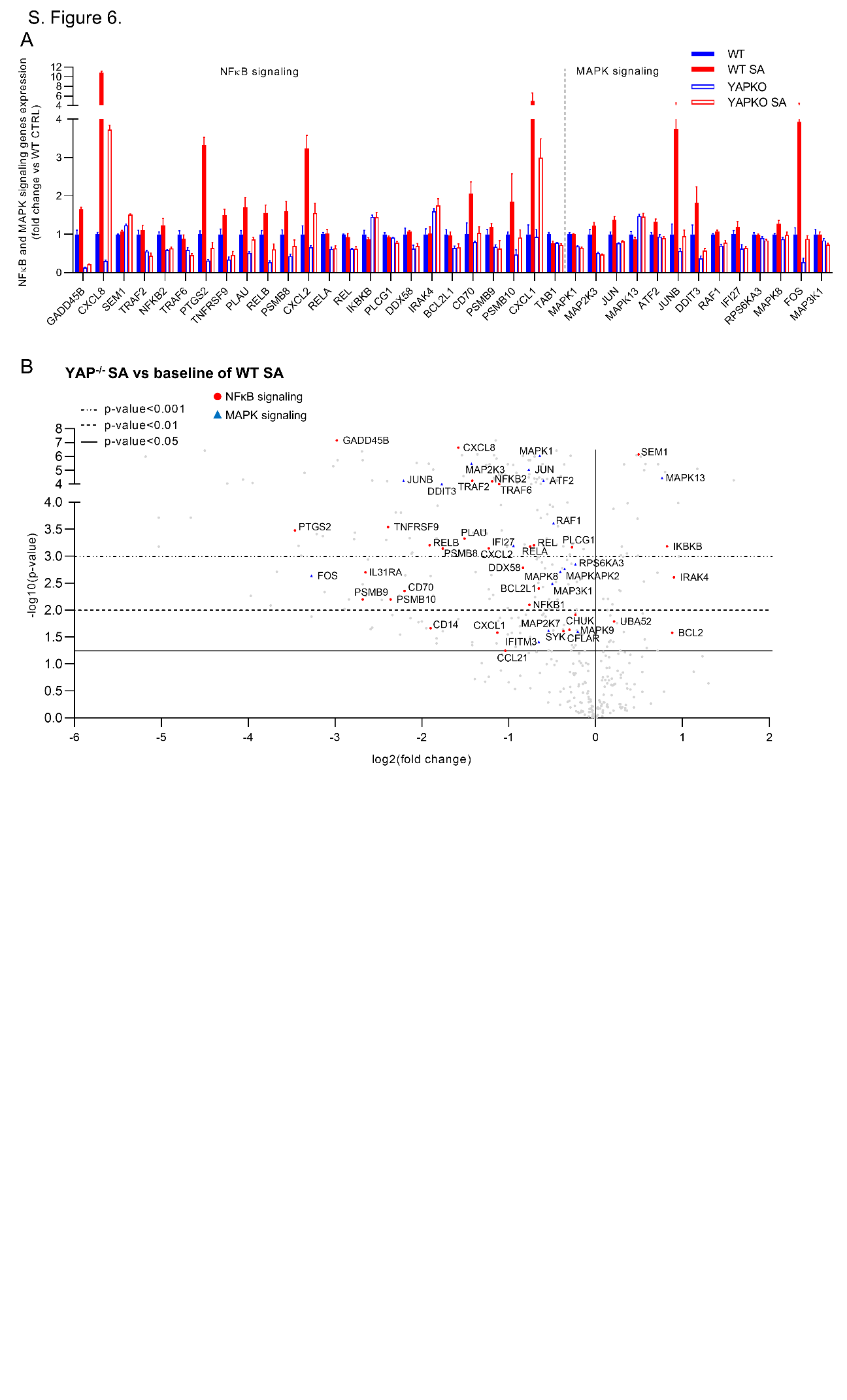
